# Propagation of information along the cortical hierarchy as a function of attention while reading and listening to stories

**DOI:** 10.1101/291526

**Authors:** Mor Regev, Erez Simony, Katherine Lee, Kean Ming Tan, Janice Chen, Uri Hasson

## Abstract

How does attention route information from sensory to high-order areas as a function of task, within the relatively fixed topology of the brain? In this study, participants were simultaneously presented with two unrelated stories – one spoken and one written – and asked to attend one while ignoring the other. We used fMRI and a novel inter-subject correlation analysis to track the spread of information along the processing hierarchy as a function of task. Processing the unattended spoken (written) information was confined to auditory (visual) cortices. In contrast, attending to the spoken (written) story enhanced the stimulus-selective responses in early sensory regions and allowed it to spread into higher-order areas. Surprisingly, we found that the story-specific spoken (written) responses for the attended story also reached the opposite secondary visual (auditory) regions. These results demonstrate how attention enhances the processing of attended input and allows it to propagate across brain areas.

## Introduction

Imagine Shibuya Crossing, the busiest junction in Tokyo. Using a single infrastructure, a dynamical traffic-control system can efficiently route traffic by switching between times that pedestrians or drivers are allowed to use the junction. Similarly, on a relatively stable long-range axonal wiring infrastructure, attentional system can control and direct the flow of incoming information based on the current task and context. For example, while reading a book at a busy coffee shop, the visual information should be routed to linguistic and extra-linguistic areas, while incoming sounds from nearby conversations that arrive at the auditory system should be suppressed. Conversely, during a phone conversation, spoken information should be routed to high-order linguistic and extra-linguistic areas, while incoming written information arriving at the visual system should be suppressed.

Selective attention can dramatically affect behavior, and therefore is likely to be associated with changes in the routing of information between early sensory and higher-order brain regions. Previous findings have consistently suggested that early sensory regions process both the attended and unattended information, but preferentially represent the attended information (Treisman, 1986; Wood and Cowan, 1995; Kastner et al., 1998; Chee et al., 1999; O’Craven et al., 1999; Jancke et al., 2001; Barrett et al., 2003; Liu et al., 2003; Golumbic et al., 2013). What is not well understood is the extent to which unattended information is processed in higher-order brain regions and the mechanism by which attended stimuli is routed from early sensory to higher-order areas.

To address these questions, we presented participants with two unrelated stories simultaneously, one spoken and one written while they underwent functional magnetic resonance imaging (fMRI; Figure 1). The two stories were unrelated in content – the spoken story depicted a tenant’s fight with an irrational superintendent, and the written story recounted a man’s personal quest to reach space. The words of the written story were presented individually and serially in the center of the screen, independent from the rhythm of the spoken words. Half of the subjects were instructed to attend the written story and ignore the spoken one (group S<u>W</u>), and the other half were asked to do the reverse (group <u>S</u>W). To ensure similar exposure to the visual content across the two groups, participants attending the spoken story were instructed to fixate their gaze on a small red dot at the center of the screen, overlapping with, but not obscuring the written words for the entire duration of the experiment. A similar design was used to map cortical areas that support the processing of attended stories irrespective of their linguistic modality (Wang and He, 2014). In this study, altered focus of attention towards written and spoken stories was used to explore how information is dynamically routed across brain regions as a function of the current cognitive task. To characterize how information is routed across brain regions as a function of task, an additional set of participants were presented with only one of the stories, constituting unimodal control groups of readers exposed only to the written story (<u>W</u>) and listeners exposed only to the spoken story (<u>S</u>). Next, we used inter-subject functional correlation (ISFC), a novel method we have developed that isolates stimulus-locked neural responses (Simony et al., 2016).

**Figure 1.**
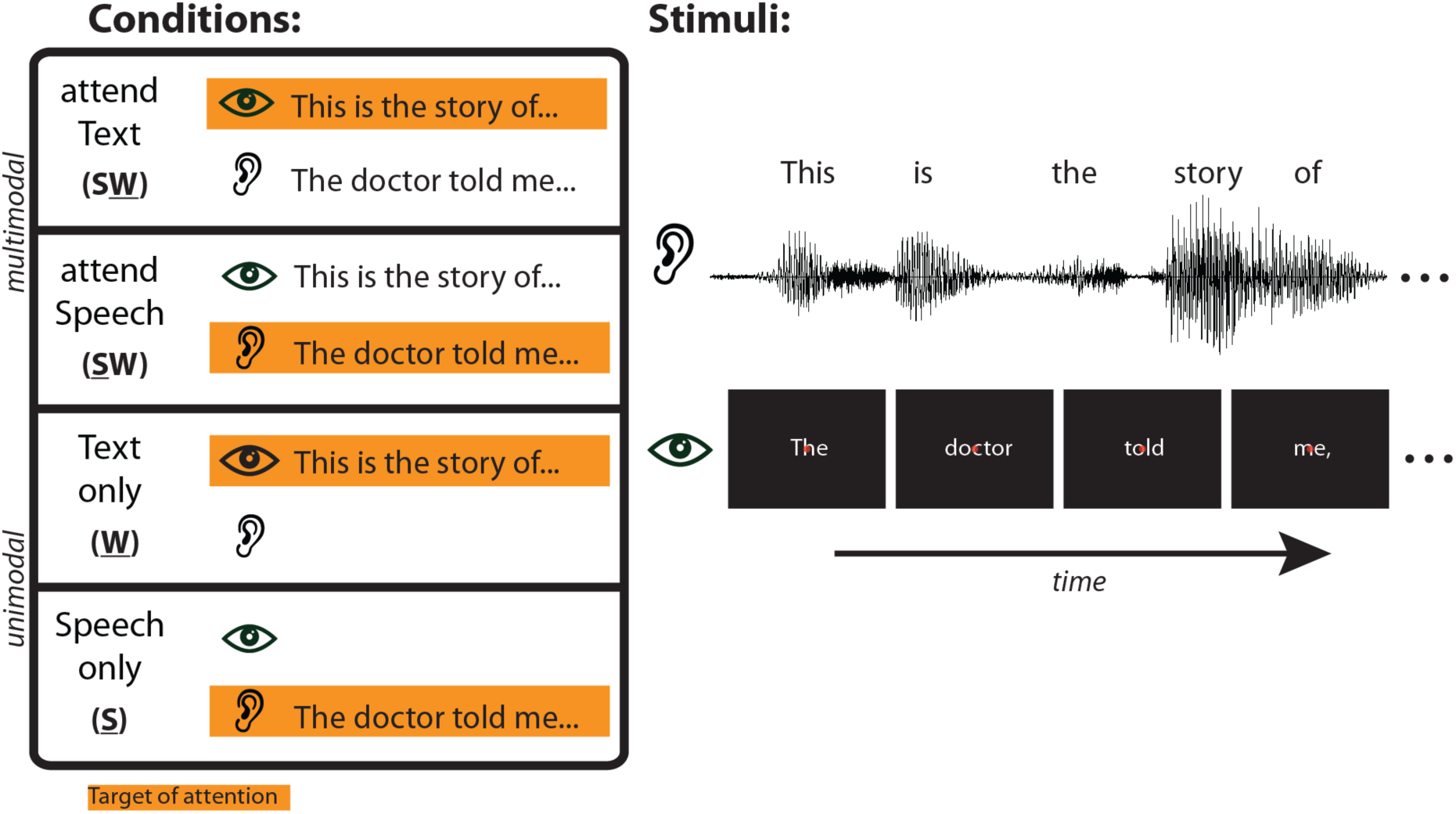
Experimental design and a short segment of each of the two 15 minute narrative stimuli. While undergoing fMRI, subjects were exposed to either one or two unrelated stories: a written story (“The Overview Effect”) and a spoken story (“Slumlord”). In the two multimodal conditions, subjects were exposed to two stories simultaneously; one group was instructed to attend only to the written story (S<u>W</u>), while the other group was instructed to attend only to the spoken story (<u>S</u>W). The two unimodal groups were exposed to and attended to only one of the stories – the spoken one (<u>S</u>) or the written one (<u>W</u>).

Inter-regional correlations measured within the same brain, such as in functional connectivity (FC) analysis, can be modeled as the sum of three components: 1) non-neuronal artifacts (e.g. physiological noise) which induce correlation across long-range areas; 2) intrinsic neural fluctuations that propagate across brain areas and can be used to uncover the layout of long-range anatomical connections during rest; 3) stimulus-locked responses that propagate across brain areas during the processing of external stimulus. While challenging, few studies have managed to isolate stimulus-locked activity from other components using FC analysis (Caclin and Fonlupt, 2006; Betti et al., 2013; Gonzalez-Castillo et al., 2015; Spadone et al., 2015). Recently, by measuring the correlation across brains (ISFC), instead of within a brain as in FC, we were able to filter out the intrinsic neural correlations and non-neural confounds (which are idiosyncratic to each brain), thus increasing the signal-to-noise ratio for detecting inter-regional correlation induced by a shared external stimulus across participants (Simony et al., 2016).

Isolating the stimulus-induced signal, while necessary, is not sufficient for capturing how attention modifies the routing of co-occurring streams of information across brain areas. Therefore, we modified ISFC to enable specific mapping of spoken and written information shared across brain areas as a function of attention. To that end, we first characterized the canonical stimulus-locked response timecourses to the spoken story and written story when presented in isolation in the two unimodal control groups. These neural responses are of great interest as they are associated with the typical attention-based processing of the narratives in daily life context. We next compared the unimodal story-specific response (<u>S</u> or <u>W</u>) to the response timecourses of the multimodal groups (<u>S</u>W and S<u>W</u>). The route of the attended spoken information was mapped by correlating the responses from the unimodal spoken story (<u>S</u>) with the responses of the multimodal group that attended the spoken story (<u>S</u>W). Similarly, the route of the attended written information was mapped by correlating the responses from the unimodal written story (<u>W</u>) with the responses of the multimodal group that attended the written story (S<u>W</u>). Conversely, the route of the unattended written information was mapped by correlating the responses from the unimodal-written story (<u>W</u>) with the responses of the multimodal group that attended the competing spoken story (<u>S</u>W), or by correlating the responses from the unimodal-spoken story (<u>S</u>) to the responses of the group that attended to competing written story (S<u>W</u>).

At the behavioral level, we found that subjects had high comprehension and recall for the attended story, and minimal comprehension and recall for the unattended stories. At the neural level, we observed that processing of the ignored spoken or written information was confined to early auditory and visual areas, respectively, and was mostly not routed further into higher-order areas. In contrast, information regarding the attended spoken or written information spread to linguistic and extra-linguistic areas, such as the angular gyrus, posterior cingulate cortex and the prefrontal cortex. Interestingly, we observed that when attention was directed toward the spoken story, secondary visual areas evidenced a considerable degree of information related to spoken story, perhaps due to top-down inter-regional interactions with higher-order executive and attention areas. Conversely, when attention was directed towards the written story, secondary auditory areas evidenced a considerable degree of information related to the written story. These results demonstrate a flexible routing of information across fixed neural networks to suit the current internal goals and demonstrate the extensive role of top-down attention in processing of naturalistic spoken and written content.

## Results

### Behavioral Results

All participants were informed, prior to attending the story, that they would later undergo a memory test about that attended story and would receive a monetary bonus based on their performance. Following the scan, participants in all groups (unimodal and multimodal) completed a questionnaire aimed at assessing their comprehension and memory for the attended story, which consisted of three tests: (a) forced choice comprehension and memory test; (b) free recall test; and (c) fill-in-the-blank test. In addition, participants in the multimodal groups were given an unexpected questionnaire about the unattended story, which they had been instructed to ignore during the scan. To motivate participants to answer the surprise questionnaire to the best of their abilities, we offered them an additional comparable monetary reward for providing correct answers for the unattended story.

Performance levels for the attended stories were high, irrespective of the presence or absence of a simultaneous second story (Fig. 2). For the spoken story, performance on all tests was similar between group <u>S</u>W (Multi: M = 84% SD = 6; Fill-in: M = 45% SD = 8; Recall: M = 9.6 SD = 5) and the control group <u>S</u> (Multi: M = 83% SD = 7.1; Fill-in: M = 44% SD = 12; Recall: M = 12, SD = 2.5; Multi: *t*_(34)_ = 0.41, *p* = 0.69; Fill-in: *t*_(34)_ = 0.4, *p* = 0.66; Recall: *t*_(33)_ = −1.67, *p* = 0.1). For the written story, performance on most of tests was similar between group S<u>W</u> (Multi: M = 87% SD = 6.2; Fill-in: M = 47% SD = 12; Recall: M = 11.4, SD = 3.2) and the control group <u>W</u> (Multi: M = 81% SD = 5.9; Fill-in: M = 49% SD = 11; Recall: M = 13 SD = 3.1; Multi: *t*_(34)_ = 2.97, *p* = 0.005, *d* = 0.39; Fill-in: *t*_(34)_ = −0.36, *p* = 0.72; Recall: *t*_(34)_ = −1.52, *p* = 0.14).

At the same time, performance levels for the unattended stories were minimal and significantly lower than the attended stories (Fig. 2). In the multiple-choice test, participants who ignored the spoken (written) story answered correctly 32%, SD = 14.2 (M = 31%, SD = 10.3) of the questions, when chance level was 25%. In the free recall test, participants who ignored the spoken (written) story scored 0.7, SD = 1 (M = 0.8, SD = 0.8) on a scale from 0 to 20. In the fill-in-the-blank test, participants who ignored the spoken (written) story answer correctly average 4%, SD = 4 (M = 11%, SD = 7) of the questions. For the spoken story, the multiple-choice (*t*_(34)_ = 14.15, *p* ≪ 0.0001, *d* = 4.9), fill-in-the-blank (*t*_(34)_ = 19.31, *p* ≪ 0.0001, *d* = 6.6), and free recall (*t*_(33)_ = 7.43, *p* ≪ 0.0001, *d* = 2.6) scores were higher in groups <u>S</u>W than in group S<u>W</u>. For written story, the multiple-choice (*t*_(34)_ = 19.9, *p* ≪ 0.0001, *d* = 6.8), fill-in-the-blank (*t*_(34)_ = 11.05, *p* ≪ 0.0001, *d* = 3.8), and free recall (*t*_(34)_ = 13.5, *p* ≪ 0.0001, *d* = 4.6) scores were higher in groups S<u>W</u> than in group <u>S</u>W.

**Figure 2.**
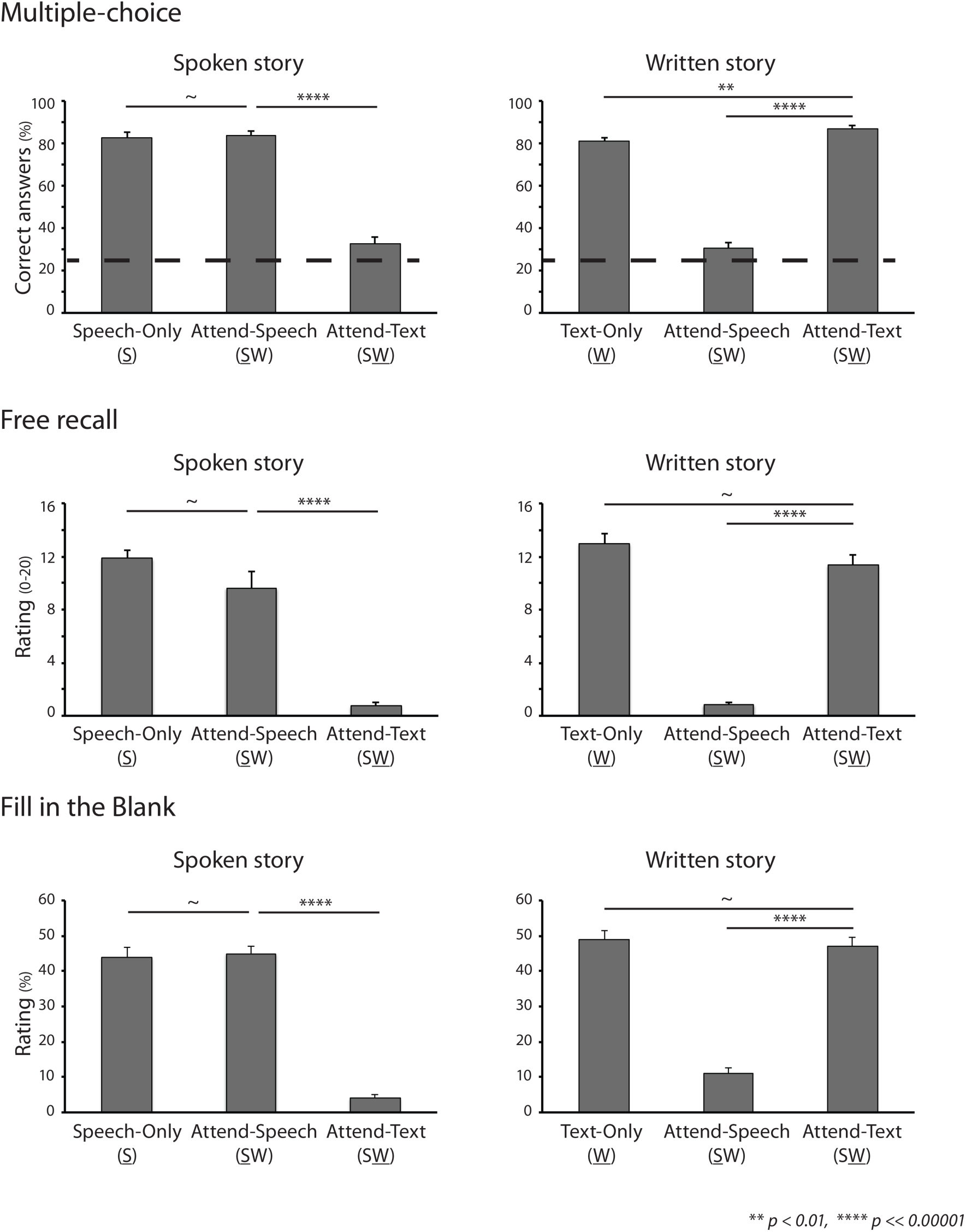
Comprehension and memory for the spoken and written stories were assessed using three post scan tests: Multiple-choice (4 options), written free recall, and fill-in-the-blank. Directing attention toward one story increased narrative comprehension and long-term memory for that story, regardless of whether the other story was simultaneously presented or absent. In all three test types, performance for the unattended stories was significantly lower than for the attended stories.

Overall, these results suggest that the research design successfully manipulated attention toward the desired story and away from the story that participants were instructed to ignore. Directing attention toward a story was associated with cognitive processing of its specific content, including increased narrative comprehension and long-term memory. In addition, the introduction of a potentially distracting story simultaneously did not seem to significantly influence participants’ comprehension and memory of the attended story. With these patterns of behavior established, we continued to explore how information was routed in the brain as a function of attention.

### Tracking task-dependent information

Initially, we characterized the neural responses to the isolated spoken and written stories in the unimodal control groups (<u>S</u> and <u>W</u>; Supplementary Fig. 1). Because the unimodal control groups were presented with only one story, their response timecourses could be treated as representing the “typical” temporal neural response patterns associate with the processing of each story in each brain region. These “typical” neural responses are of interest as they are associated with attention-based processing of the narratives and their in-depth comprehension, as validated in the behavioral test. The temporal neural responses collected from the unimodal groups were each compared to both of the multimodal groups (<u>S</u>W and S<u>W</u>), using Pearson correlation. This allowed us to model responses of each multimodal group with respect to the attended as well as the unattended stimuli. Note that this design only enables tracking of changes in the typical processing of the spoken or written stories as a function of attention, when presented in the multimodal conditions. It is not intended to capture any responses which are idiosyncratic to the multimodal mode of presentation that are not present in the typical unimodal mode of presentation.

#### Processing of unattended information

To map areas that preserved their response pattern to the incoming input, irrespective of attentional control, we compared the typical responses of a unimodal group with the responses of the multimodal group who did not attend the same story (<u>S</u> subjects with S<u>W</u> subjects, and <u>W</u> subject with <u>S</u>W subjects). When this cross-group comparison is performed using the *same* brain area in both groups, which we denote as an inter-subject correlation (ISC) analysis (Fig. 3A, thick line), we can map areas that responded in a similar way whether the story was attended or ignored. For example, by assessing whether neural responses in the auditory cortex are similar when participants ignore the presented spoken story (S<u>W</u>), compared to when they attend to it without distraction (<u>S</u>), we can identify processing of spoken information irrespective of attention within the auditory cortex. When the cross-group comparison is performed between *different* areas across the groups, which we denote as an inter-subject functional correlation (ISFC) analysis (Fig. 3A, dashed lines), we can map how the stimulus-locked responses to the unattended story are shared across two brain areas. For example, we can test whether spoken information is shared between the auditory cortex and the angular gyrus by assessing whether neural response in the auditory cortex when participants attend the competing written story (S<u>W</u>), is correlated with the response in the angular gyrus when they attend the spoken story without distraction (<u>S</u>).

**Figure 3.**
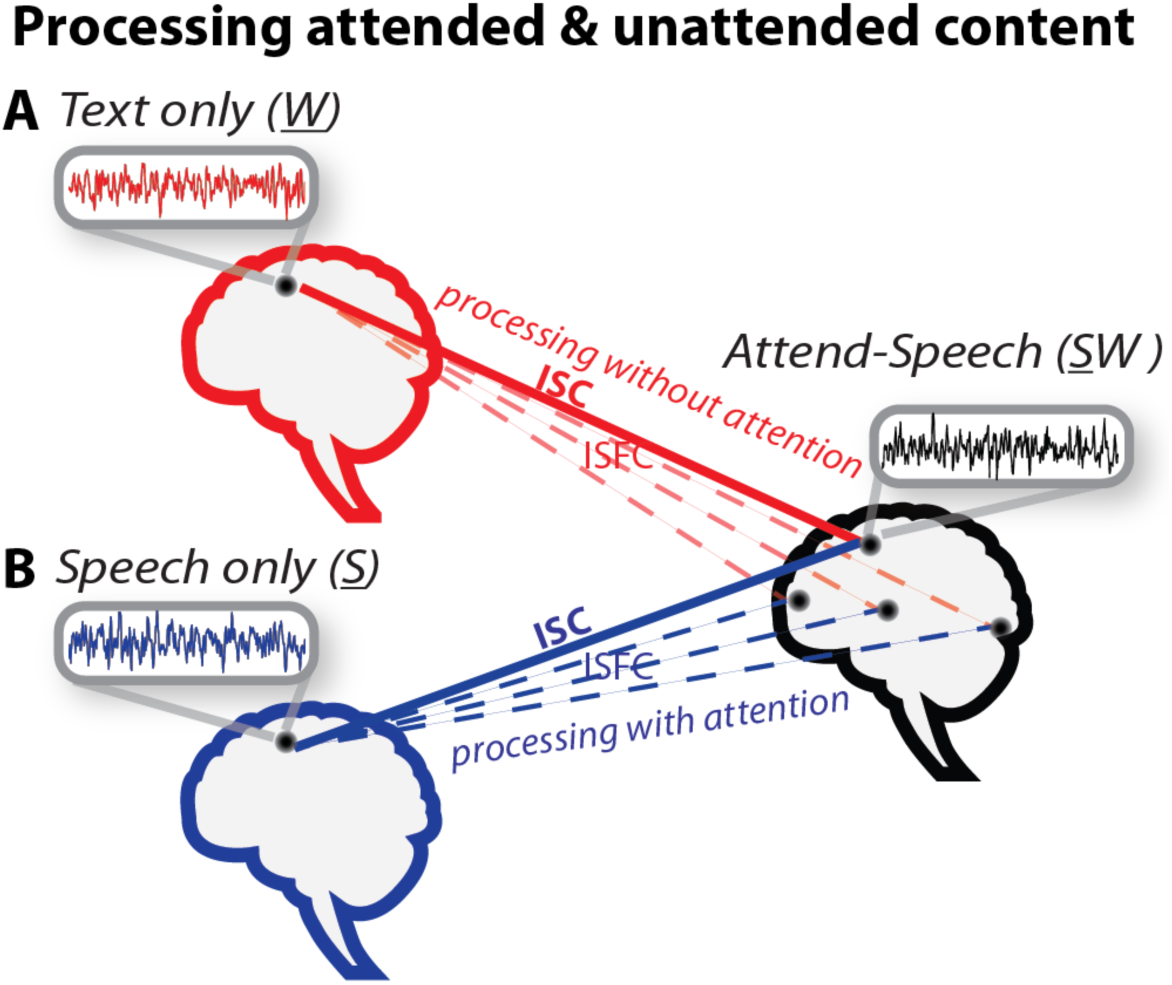
Inter-subject correlation of the BOLD timecourses was performed across groups either using the same brain areas in each group (ISC, thick lines), or between different brain areas (ISFC, dashed lines). Neural responses of the multimodal groups who attended speech or text (<u>S</u>W or S<u>W</u>) were compared with responses from the two unimodal control groups (<u>W</u> and <u>S</u>), which represent the “typical” response to the written (red) or the spoken (blue) story. **A,** Inter-subject correlations between group <u>S</u>W and group <u>W</u> reveal processing of the ignored written content (red). **B,** Inter-subject correlations between group <u>S</u>W and group <u>S</u> reveal processing of the attended spoken content (blue).

#### Processing of attended information

To map areas that preserved their response pattern while the stories were attended, we compared the typical responses of a unimodal group with the responses of the multimodal group who attended the same story (<u>S</u> subjects with <u>S</u>W subjects, and <u>W</u> subjects with S<u>W</u> subjects). When this cross-group comparison is performed using the *same* brain area in ISC analysis (Fig. 3B, thick line), we can map areas that responded similarly to the attended stories when presented in isolation compared to in the multimodal condition. Additionally, when the cross-group comparison is performed between *different* areas in ISFC analysis (Fig. 3B, dashed lines), we can map how attended information is shared across two brain areas. For example, by assessing whether the neural response in the auditory cortex when participants attend the spoken story (<u>S</u>W) is correlated with the response in the angular gyrus when that story was attend without distraction (<u>S</u>), we can identify information from the spoken story that was shared across these two areas while that story is attended.

The ISC and ISFC analyses were performed at the voxel level as well as on timecourses from 61 independently defined regions of interest (ROIs), which were created out of six intrinsic connectivity networks (see Materials and Methods; see Fig. 4, Table 1; see Supplementary Fig. 2) using a parcellation approach (Baldassano et al., 2015). The correlation values (ISC and ISFC) were corrected for multiple comparisons using a bootstrapping procedure based on phase randomization (see Materials and Methods, under *ISFC and ISC bootstrapping and phase-randomization*).

**Figure 4.**
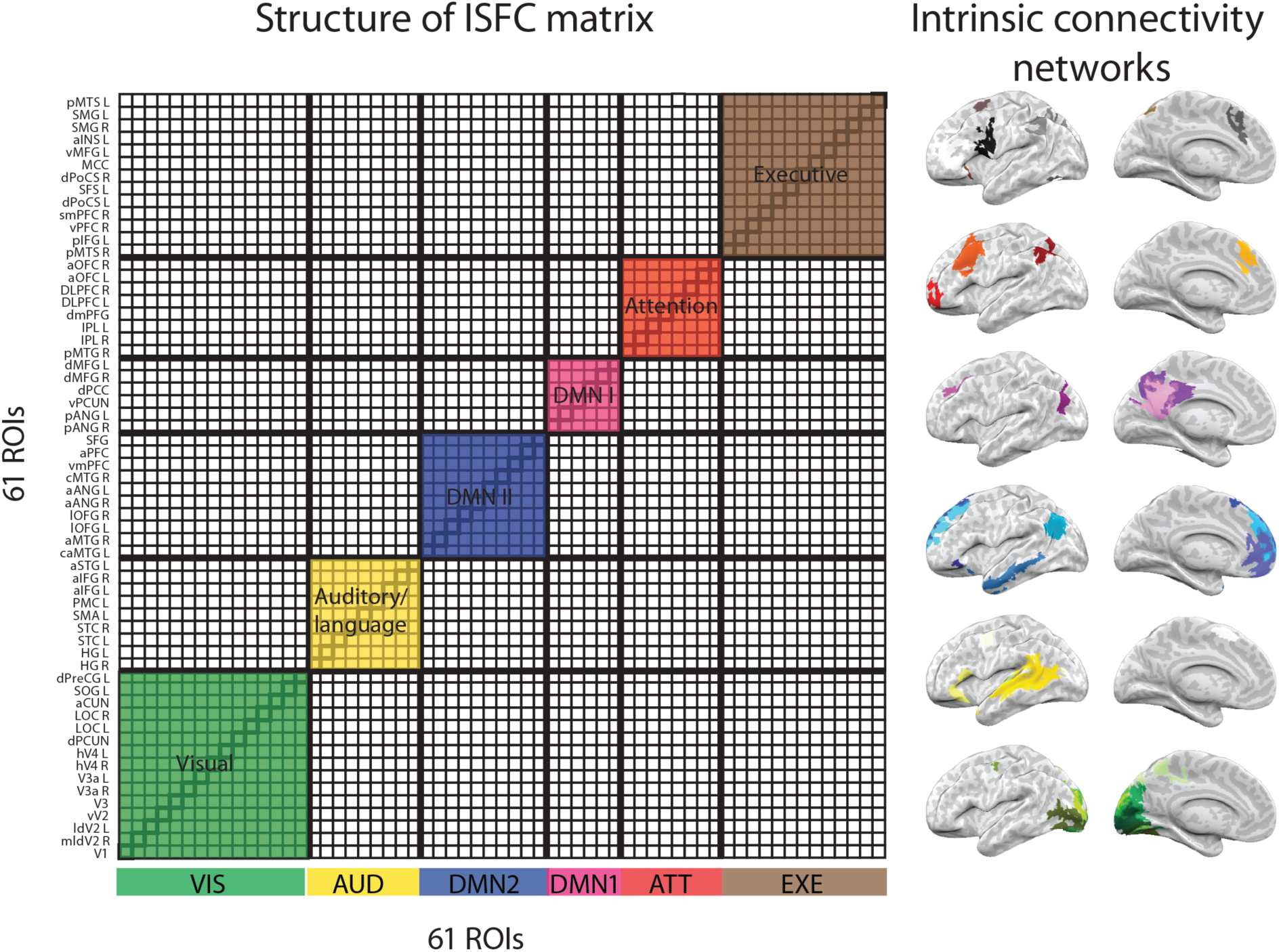
ISC and ISFC analyses were performed on time courses from 61 independently defined regions of interest (ROIs), which were created out of six intrinsic connectivity networks (ICNs) using a parcellation approach. The correlation matrix computed across brains includes an inter-subject comparison within ROIs (the diagonal represents ISC), between ROIs of the same network (colored off-diagonal), and between ROIs of different networks (uncolored off-diagonal). See Supplementary Fig. 2 for a more detailed localization of ROIs within the ICNs.

**Table 1.**
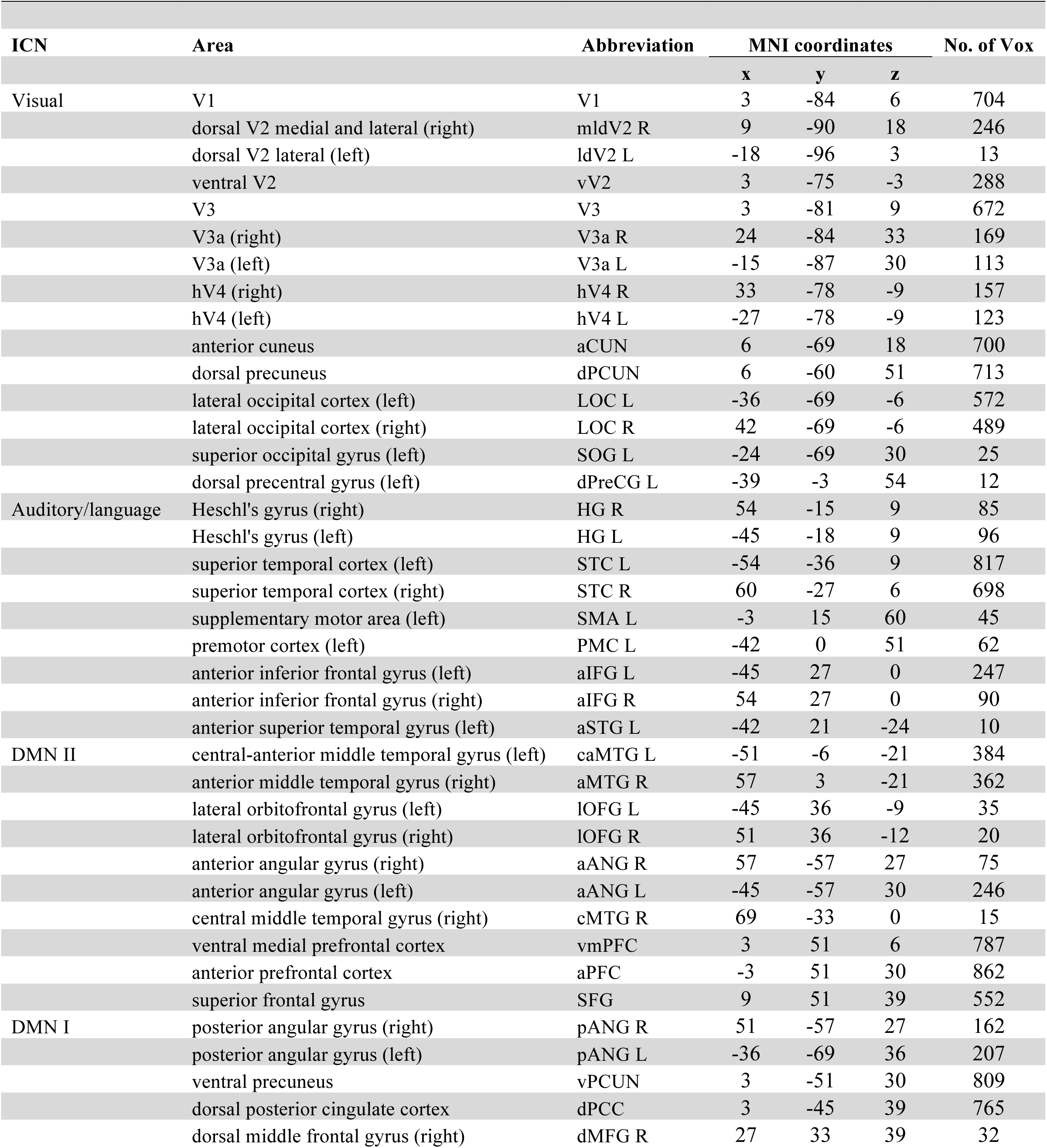
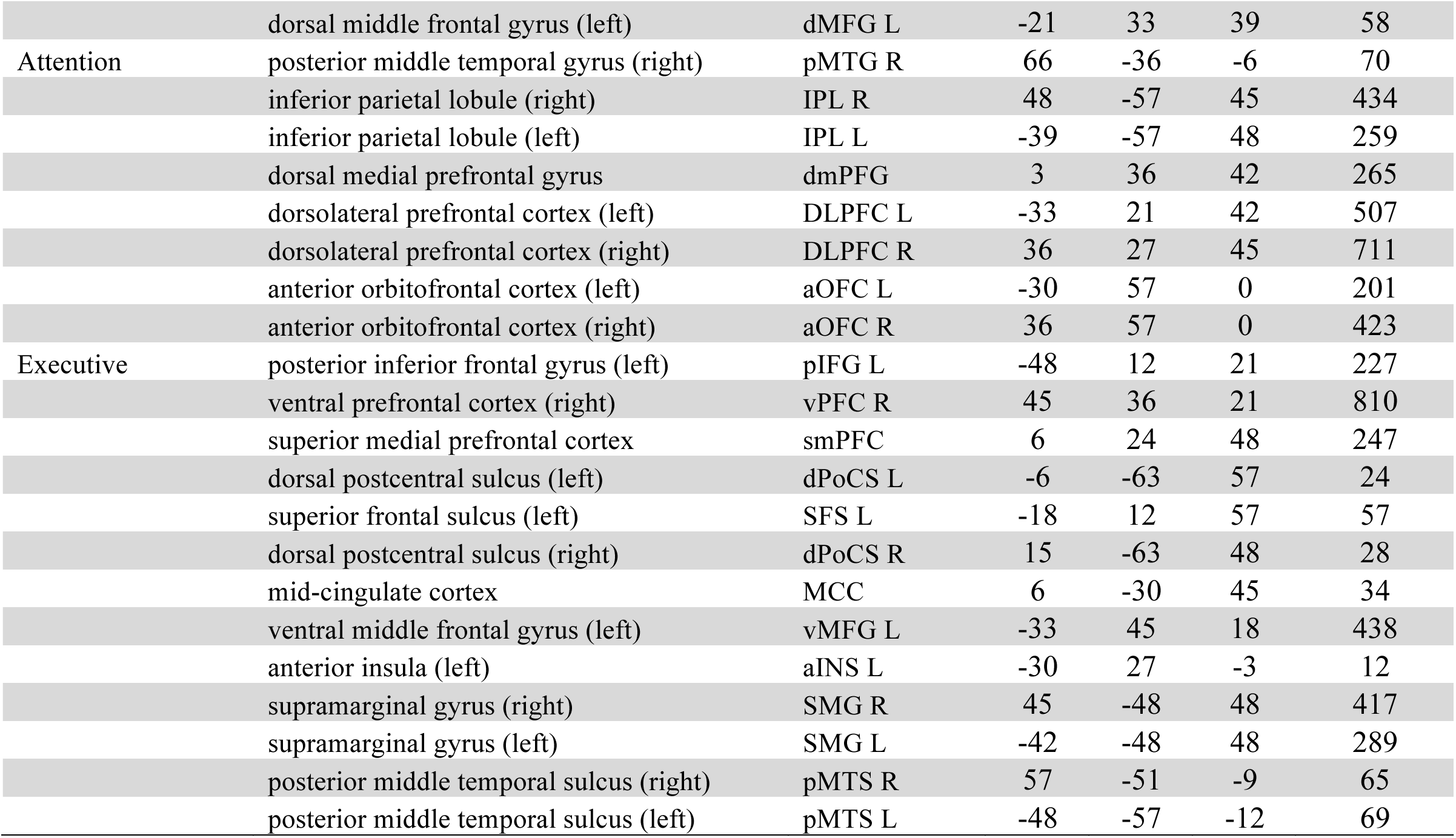
61 ROIs

### Propagation of unattended information across cortical areas

Unattended spoken and written stories evoked typical response timecourses in the modality-appropriate sensory cortices of the multimodal groups. When participants ignored the spoken story (S<u>W</u>), their response within Heschl’s gyri and nearby regions along the superior temporal cortex was similar to the typical response of the unimodal listeners (<u>S</u>; *p*_(*FWER*)_ ≪ 0.0001; Fig. 5B). Furthermore, the inter-regional correlations for the unattended spoken story seemed to be restricted mainly to auditory areas (see off diagonal in ISFC matrix in Fig. 5A, wherein a cluster of positive correlations was observed primarily among auditory areas, including Heschl’s gyri, STC, PMC L, aSTG L, caMTG L, aMTG R, c MTG R, aIFG, and lOFG; *p*_(*FWER*)_ ≪ 0.0001). Interestingly, the inter-regional correlations did not propagate from the auditory cortex to higher order linguistic and extra-linguistic areas.

**Figure 5.**
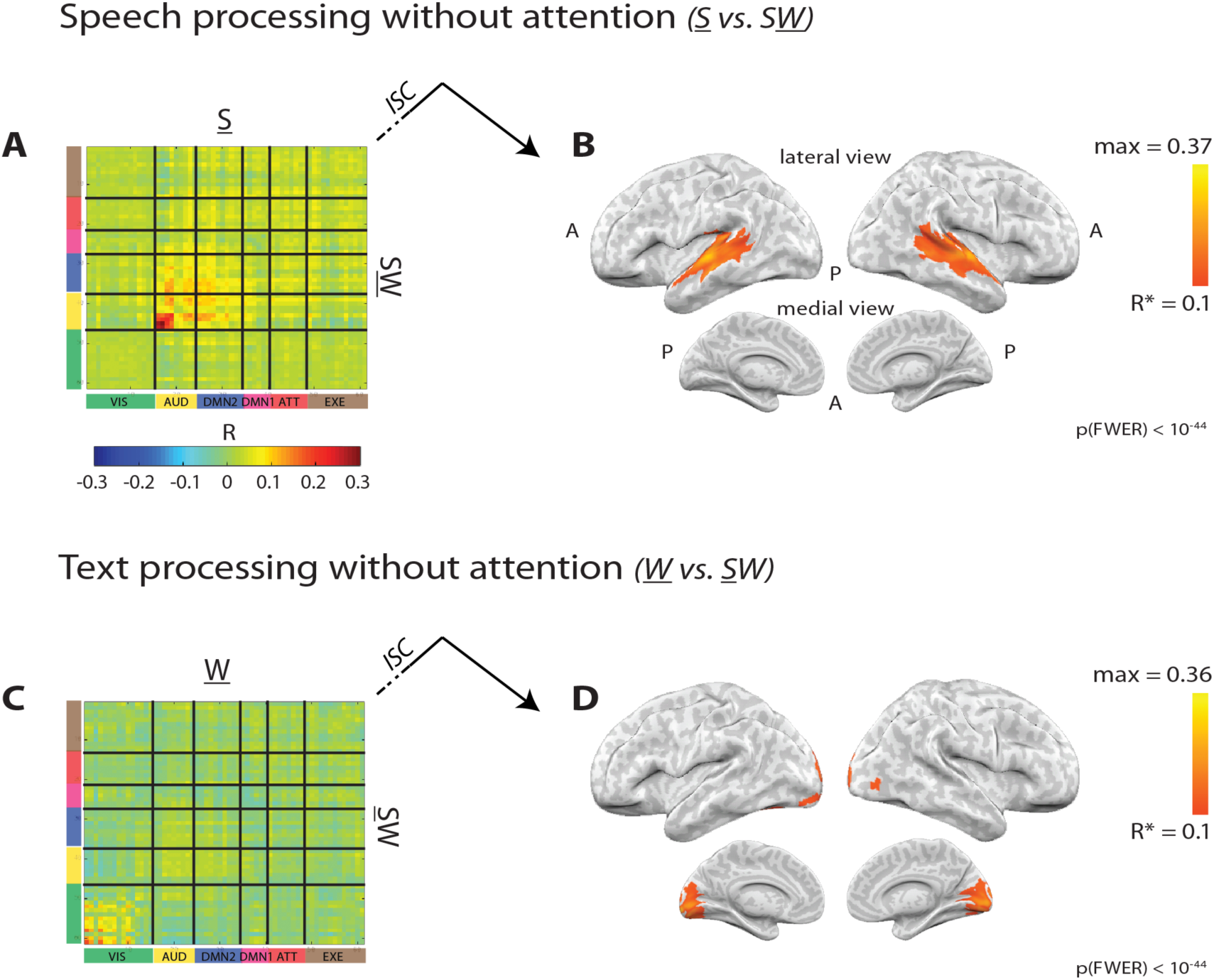
Unattended stories evoked typical processing responses in sensory cortices. **A,** When the spoken story was ignored (multimodal group S<u>W</u>) an ROI-based ISFC analysis showed that a typical (i.e., highly similar to the <u>S</u> unimodal group) response to the spoken story was shared between auditory regions. **B,** A voxel-based ISC analysis showed a typical response to the ignored spoken story within auditory regions (*p*_(6789)_ < 10^−44^). **C,** When the written story was ignored (multimodal group <u>S</u>W) an ROI-based ISFC analysis showed a typical (i.e. highly similar to the <u>W</u> unimodal group) response to the written story shared between visual regions. **D,** A voxel-based ISC analysis showed a typical response to the ignored written story within visual regions (*p*_(6789)_ < 10^−44^).

Similarly, when participants ignored the written story (<u>S</u>W), their response was similar to the typical response of the unimodal readers (<u>W</u>) within the visual system, including V1, V2, V3, V3a, hV4, aCUN, and LOC (*p*_(*FWER*)_ ≪ 0.0001; Fig. 5D). A restricted inter-regional correlation was observed here as well, wherein a cluster of inter-region correlations was observed primarily among visual areas (see off diagonal in ISFC matrix in Fig. 5C). These regions included V1, mldV2 R, vV2, V3a L, and aCUN (*p*_(*FWER*)_ ≪ 0.0001). This result of inter-regional responses being restricted to sensory cortices were reproduced when the analysis was performed at the voxel level, instead of the ROI level (see Supplementary Fig. 3C and D), and when mutual dependencies between regions were removed (see Materials and Methods), to reflect more direct inter-regional interactions (i.e. partial correlation; Supplementary Fig. 4C and D).

### Propagation of attended information across cortical areas

Attention to speech allowed the information from the spoken story to propagate from auditory areas to higher-order regions (Fig. 6A and B). In marked contrast to the unattended condition (S<u>W</u>; Fig. 5B), attending to the spoken information (<u>S</u>W) induced responses which were highly correlated with the responses seen when processing the spoken story in isolation (<u>S</u>) in linguistic and extra-linguistic areas (*p*_(*FWER*)_ ≪ 0.0001; Fig. 6B). Furthermore, the spoken information seemed to propagate across many of the nodes within and between the Attention network, Executive network, and Default Mode Networks (DMNs; see off diagonal in ISFC matrix in Fig. 6A). In particular, we observed increased correlation between auditory and linguistic areas, between linguistic areas and the DMNs, and between the Attention and Executive networks (*p*_(*FWER*)_ ≪ 0.0001).

**Figure 6.**
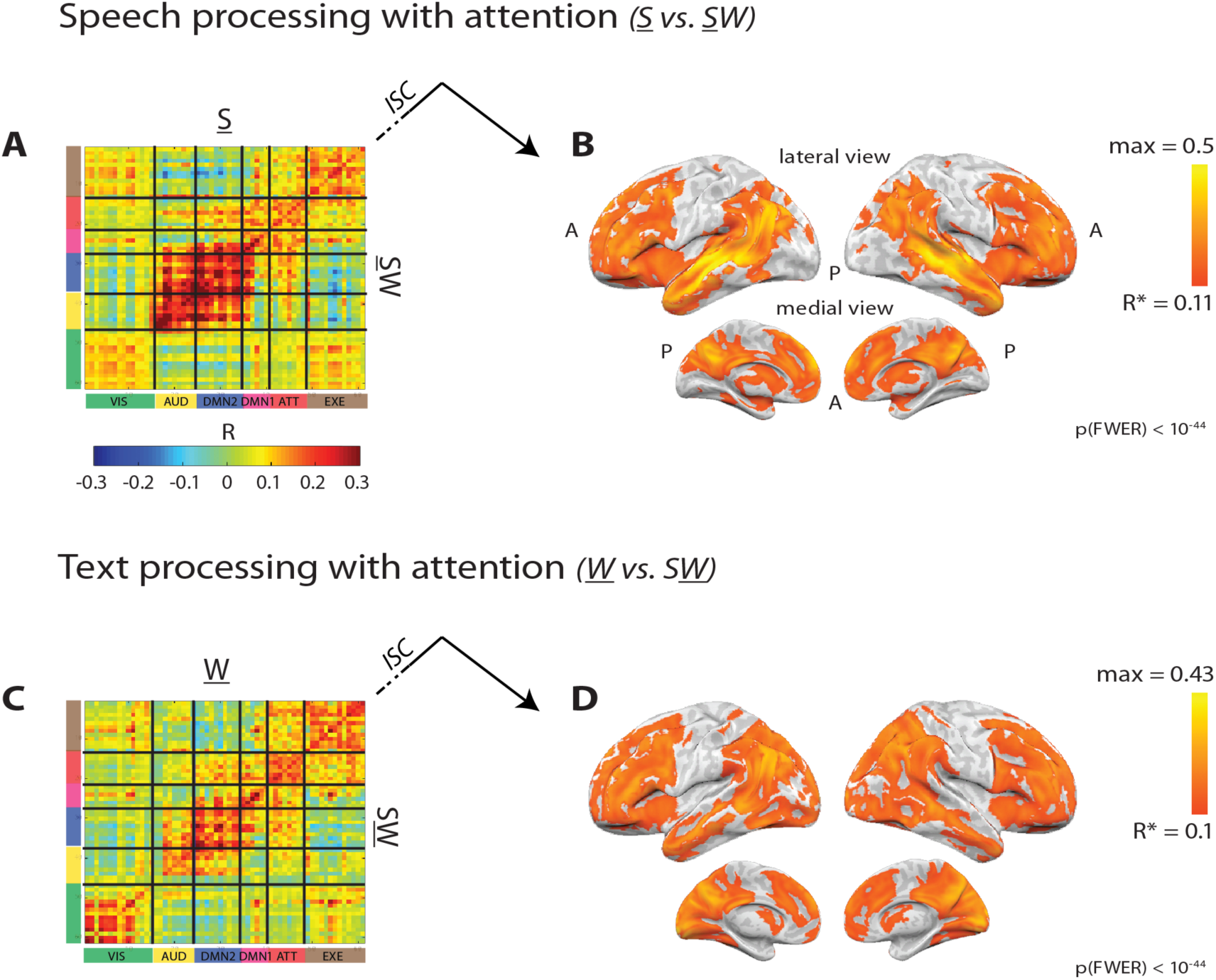
Attention to stories evoked typical processing responses that spread from sensory regions to high-order areas. **A,** When the spoken story was attended (multimodal group <u>S</u>W) an ROI-based ISFC analysis showed that a typical response to the spoken story was shared between both auditory cortices and high-order areas. **B,** A voxel-based ISC analysis showed a typical response to the attended spoken story within auditory and high-order areas (*p*_(*FWER*)_ < 10^−44^). **C,** When the written story was attended (multimodal group S<u>W</u>) an ROI-based ISFC analysis showed that a typical response to the written story was shared between both visual cortices and high-order areas. **D,** A voxel-based ISC analysis showed a typical response to the attended written story within visual and high-order areas (*p*_(*FWER*)_ < 10^−44^).

Similarly, attention to text modified the spread of information from the written story, allowing it to propagate from early visual to higher-order areas (Fig. 6C and D). The responses to the attended written story (S<u>W</u>) were highly correlated with the responses to the written story in isolation (<u>W</u>) in linguistic and extra-linguistic areas (*p*_(*FWER*)_ ≪ 0.0001; Fig. 6D). Furthermore, in the S<u>W</u> condition the written information (but not the ignored spoken information) seemed to propagate across many of the nodes within and between the DMNs, Attention network and the Executive network (see off diagonal in ISFC matrix in Fig. 6C). In particular, we observed increased functional correlation between high-order visual cortices (especially the lateral occipital cortex, superior occipital gyrus and the anterior cuneus) with regions in the Executive network and DMN-I (*p*_(*FWER*)_ ≪ 0.0001). These inter-regional correlations were reproduced when the analyses were performed on the voxel level, instead of the ROI level (see Supplementary Fig. 3A and B), and when mutual dependencies between regions were removed (see Supplementary Fig. 4A and B).

Overall, when participants were exposed to the two stories simultaneously through two different modalities, attention toward one modality enhanced the neural response of story-specific information and allowed it to propagate from early sensory regions to high-order linguistic and extra-linguistic areas. In other words, when subjects attended the spoken story, there was little to no trace of the written story information in linguistic and extra-linguistic areas (see empty map in <u>W</u> vs. <u>S</u>W comparison, Fig. 5C and D). Instead, these regions were dominated by shared spoken information (see correlations in <u>S</u> vs. <u>S</u>W comparison, Fig. 6A and B). However, attention to the written story reversed this effect: responses associated with the written story were now found in high-order regions (see correlations in <u>W</u> vs. S<u>W</u> comparison, Fig. 6C-D), and traces of spoken information in these areas were diminished (see empty map in <u>S</u> vs. S<u>W</u> comparison, Fig. 5A and B). At the same time, information from the simultaneously presented but unattended written (or spoken) story was evident mainly within the early visual (or auditory) cortices.

Notably, this analysis could only detect response patterns that are similar to the typical responses observed when the spoken or written stories were presented in isolation and fully comprehended. The neural responses that are distinctive to the simultaneous multimodal presentation of the stories remains to be characterized in future studies.

### Inter- and intra-regional modulation by attention

Previous work has shown that the degree to which attention enhances neural responses varies across regions along the processing hierarchy (Jancke et al., 2001; Bluvas and Gentner, 2013; Golumbic et al., 2013; Wang and He, 2014). To describe the degree of neural response enhancement by attention, we calculated a second order *Attention-Index* (*AI*) that assessed the difference in response for the same spoken or written story when it was attended versus unattended (for more details see Materials and Methods, under *Attention Index*). The more positive the *AI* is, the more reliable the neural response while attended, while negative *AI* values indicate that the responses are more reliable without attention, and an *AI* value close to zero indicates that the neural responses to the stories is similar for the attended and unattended stories.

Although visual and auditory regions responded reliably to text and speech (respectively) even when ignored, the reliability of responses were somewhat enhanced when the stories were fully attended. Attention increased the reliability of responses and inter-regional correlations in sensory regions by up to 33%, as reflected by the positive *AI* values (see yellow areas in Fig. 7A-D). In higher-order brain regions, which include the Attention, Executive, and Default Mode Networks, the *AI* consistently showed a sharper increase in response reliability of about 33-100% (see red areas in Fig. 7A-D). The high *AI* values in high-order regions are a result of weak reliability and inter-regional correlation of unattended information (Fig. 5). To assess the significance of the enhancement in response reliability by attention in different cortical areas, the reliability of response (ISC) was compared within a sample of sensory and high-order regions of interest, using a *t* test (see Materials and Methods). A significant enhancement in reliability by attention was detected in both high-order as well as early sensory regions (Fig. 7E). Response reliability was strongly enhanced when the stories were attended in linguistic and extra-linguistic regions such as the STC L, aANG L and dPCC (*p* ≪ 0.0001, *d* > 1.67), but also in V1 and Heschl’s gyrus (*p* ≪ 0.0001, *d* > 0.8).

**Figure 7.**
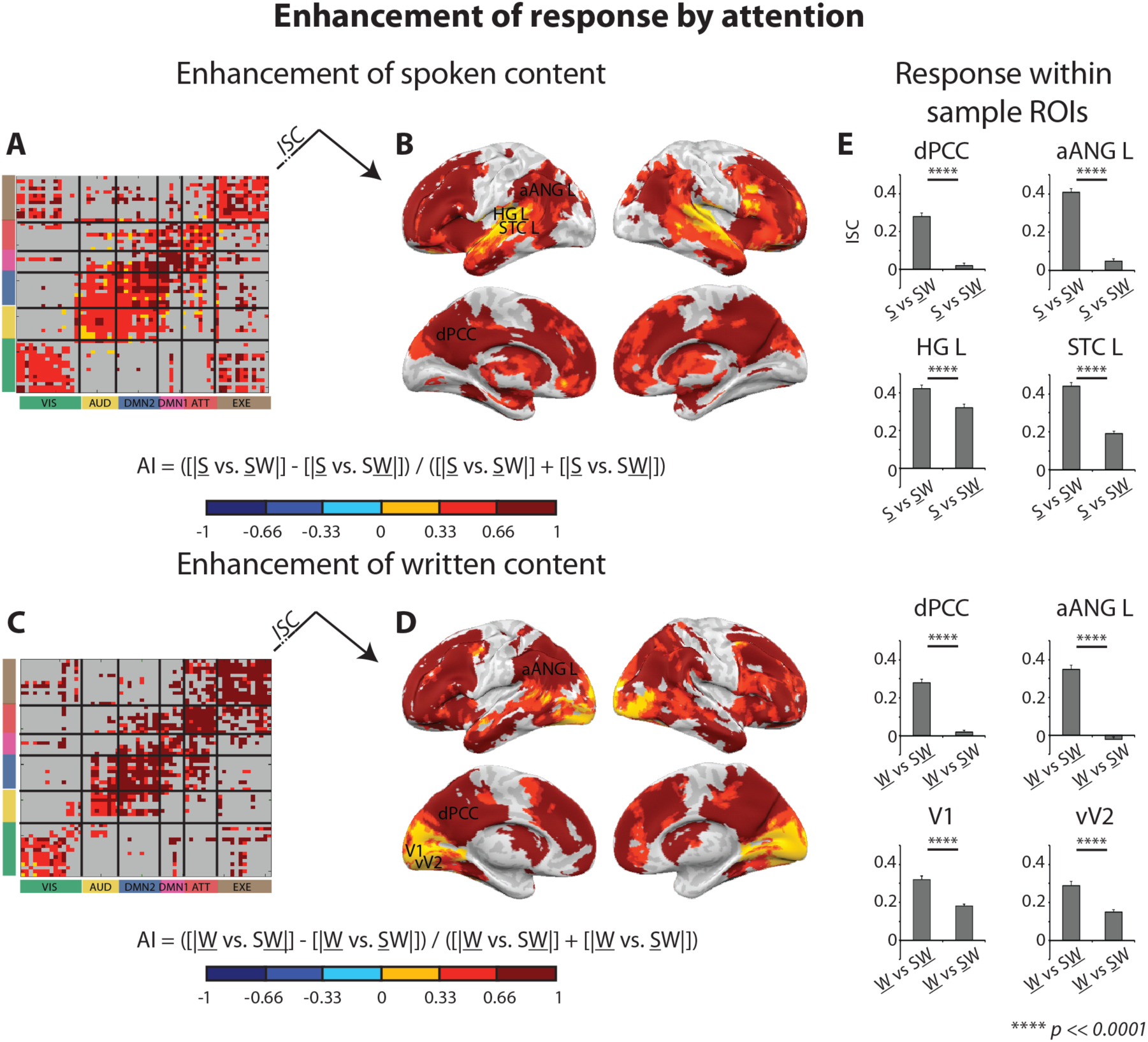
Attention to stories enhanced inter- and intra-regional responses differently along the processing hierarchy. Degree of enhancement was assessed by calculating a second order attention-index (AI) of the difference in response to the story in its attended and unattended states, in proportion to the sum of responses in both states. Cortical areas that did not show significant response to either of the attentional states were excluded from this analysis (*p*_(*FWER*)_ ≪ 0.0001; grey colors). When the spoken story was attended (<u>S</u>W), inter-regional (**A**) and intra-regional (**B**) responses to the spoken story were strongly enhanced in high-order ROIs (red colors), but weakly enhanced mostly in auditory cortices (yellow color). When the written story was attended (S<u>W</u>), inter-regional (**C**) and intra-regional (**D**) responses to the written story were strongly enhanced in high-order ROIs, but weakly enhanced in early visual cortices. **E,** Responses to stories in high-order and sensory ROIs (dPCC, aANG L, STC L, vV2 and V1) were stronger when they were attended vs. when ignored.

### Inter- and intra-regional segregation of information

The detection of both attended and unattended information simultaneously in the brain raises the question of how mixed the response patterns are to the two stimuli in each brain area. In one extreme case, the stimuli might be completely segregated spatially in the brain, with only the attended or unattended signals present in each brain area. This would suggest that attention fully blocks routing of unattended information into areas that are involved in processing the attended input stream, and/or blocks attended information from reaching areas that process the unattended content. In the opposite extreme case, we may find that some regions equally response to both the attended and unattended stories, showing no spatial segregation in the processing of relevant and irrelevant content.

In the following analysis, we measured the extent to which each region maintained both attended and unattended signals simultaneously. We calculated a second order *Segregation-Index* (*SI*) which compared ISC or ISFC levels between the simultaneously presented attended and unattended stories (for more details see Materials and Methods, under *Segregation Index*). According to this descriptive index, areas that primarily process the attended story would show an *SI* value close to one, while areas that primarily process the unattended content would show an *SI* value close to minus one, and areas that process the attended and unattended stimuli to an equal extent would have an *SI* value close to zero.

Segregation levels between the attended and unattended stories varied along the cortical processing hierarchy. Responses in high-order regions were dominated by the attended stories, with little trace of response to the unattended content (Fig. 8, red colors, SI close to 1). Responses in early auditory areas (left and right Heschl’s gyri), and to lesser extent in primary visual cortex, were found to be mainly dominated by the modality-appropriate sensory input even when it was not attended (Fig. 8, blue colors, SI close to −1). Unexpectedly, when the spoken story was attended (<u>S</u>W), responses to the spoken story were also observed in visual cortices and mixed with responses to the unattended text (Fig. 8A and B, cyan and yellow, SI close to 0). Conversely, when the written story was attended (S<u>W</u>), responses to the written story reached high-level auditory regions (Fig. 8C and D, cyan and yellow, SI close to 0). Thus, while the visual (or auditory) areas were involved in processing the unattended sensory information, at the same time, they seemed to receive input related to the information coming from the competing auditory (or visual) stimulus. To assess the significance of the segregation between the responses in different cortical areas, the reliability of response (ISC) was compared within a sample of sensory and high-order regions of interest, using a *t* test (Fig. 8E). A strong dominance of responses to the attended story was observed in higher-order regions (aANG L: *p* ≪ 0.0001, *d* > 2.16, dPCC: *p* ≪ 0.0001, *d* > 1.98); and a strong dominance of responses to the sensory stimuli (regardless of attention) was observed in early sensory cortices (V1: *p* < 0.005, *d* > 0.51, HG L: *p* ≪ 0.0001, *d* > 2.28). Interestingly, a more balanced amount of response to the unattended and attended stories (from the opposite sensory modalities) was observed in secondary visual and auditory regions of the unattended modality (vV2 in <u>S</u>W: *p* = 0.2, *d* = 0.22, STC L in S<u>W</u>: *p* = 0.23, *d* = 0.21).

**Figure 8.**
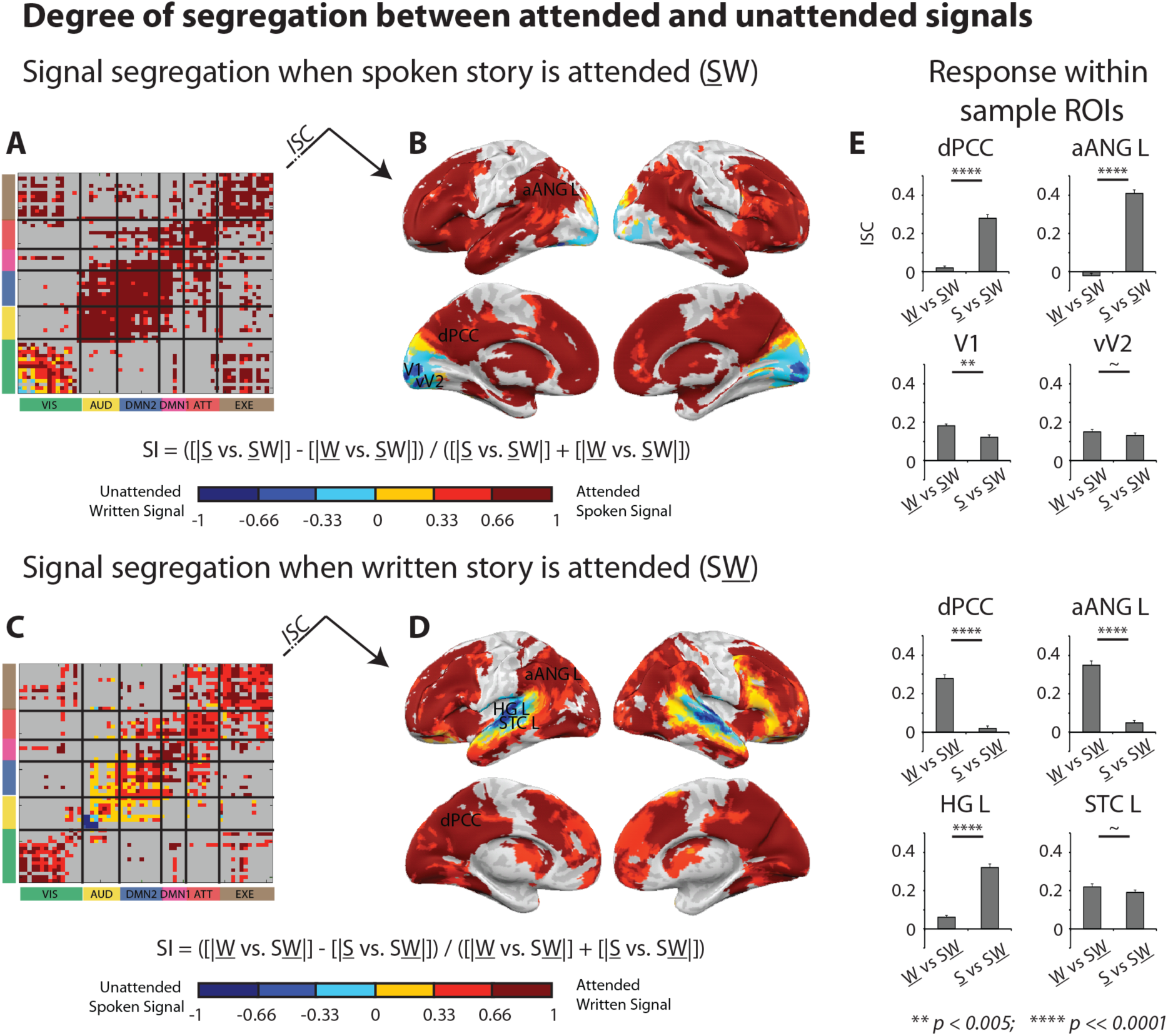
Spatial segregation between responses to the simultaneously presented stories differ along the processing hierarchy. Degree of segregation was assessed by calculating a second order segregation-index (SI) of the difference in response to the attended and unattended stories, in proportion to the sum of responses to both stories. Cortical areas that did not show significant response to either of the simultaneously presented stories were excluded from this analysis (*p*_(*FWER*)_ ≪ 0.0001; grey colors). When the spoken story was attended (<u>S</u>W), inter-regional (**A**) and intra-regional (**B**) responses to the spoken story dominated high-order areas over responses to the ignored written story (red colors). At the same time, early visual areas were dominated by inter-regional responses to the ignored written story (blue colors), while some high-order visual cortices include responses to both the spoken and the written stories (cyan and yellow). When the written story was attended (S<u>W</u>), inter-regional (**C**) and intra-regional (**D**) responses to the written story dominated high-order areas over responses to the ignored spoken story (red colors). At the same time, early auditory regions were dominated by inter-regional responses to the ignored spoken story (blue colors), while some high-level auditory cortices also include responses to the written and the spoken stories (cyan and yellow). **E,** Responses in high-order ROIs (e.g. dPCC, aANG L) to the attended story dominated over responses to the simultaneously presented but ignored story. Responses in early sensory ROIs (e.g. HG L, V1) to the story in the relevant sensory modality (whether attended or ignored) dominated over responses to the simultaneously presented story in the irrelevant modality. In secondary sensory ROIs (e.g. STC L, vV2) there were similar levels of responses to the simultaneously presented attended and ignored stories.

These results demonstrate the extent to which attention allows the relevant content to propagate across brain regions. Information related to the attended stories not only reached high-order regions, but surprisingly was also reached sensory regions that processed the unattended sensory modality.

### Input-dependent propagation of information

While information from both the written and the spoken stories reached a similar set of linguistic and extra-linguistic areas when attended, the different sensory origins of the stories compel them to undergo at least partly distinctive processing pathways (i.e. speech must go through an auditory processing pathway before reaching linguistic areas, while text must go through a visual processing pathway). To characterize alterations in the functional pathways of attended spoken versus written stories, we further examined the inter-regional correlation matrices which describe the spread of attended spoken (ISFC of <u>S</u> with <u>S</u>W) and written (ISFC of <u>W</u> with S<u>W</u> subjects) information across brain regions.

First, we searched for areas that modified their inter-regional correlation patterns as a function of the attended story (i.e. areas that were coupled with different sets of regions when speech was attended vs. when text was attended). The ISFC pattern of each ROI was represented as a vector whose angle and length can be defined relative to the origin (Wang et al., 2015). For each region, one vector depicted correlation values with other brain regions for the attended written content (<u>W</u> with S<u>W</u>) and a second vector depicted correlation values with other brain regions for the attended spoken content (<u>S</u> with <u>S</u>W). We extracted the angle change (cosine distance) between the two connectivity vectors, with large distance between the vectors capturing a large change in goal-related inter-regional correlation patterns (see Materials and Methods). The strongest changes in routing of spoken versus written content was observed in early sensory regions, but also in the pIFG L, the SMG R, and the STC R (Fig. 9). The weakest changes in routing of spoken versus written information (*cos* < 0.05) were observed mainly within regions in the prefrontal and inferior parietal cortices (see Table 2).

**Figure 9.**
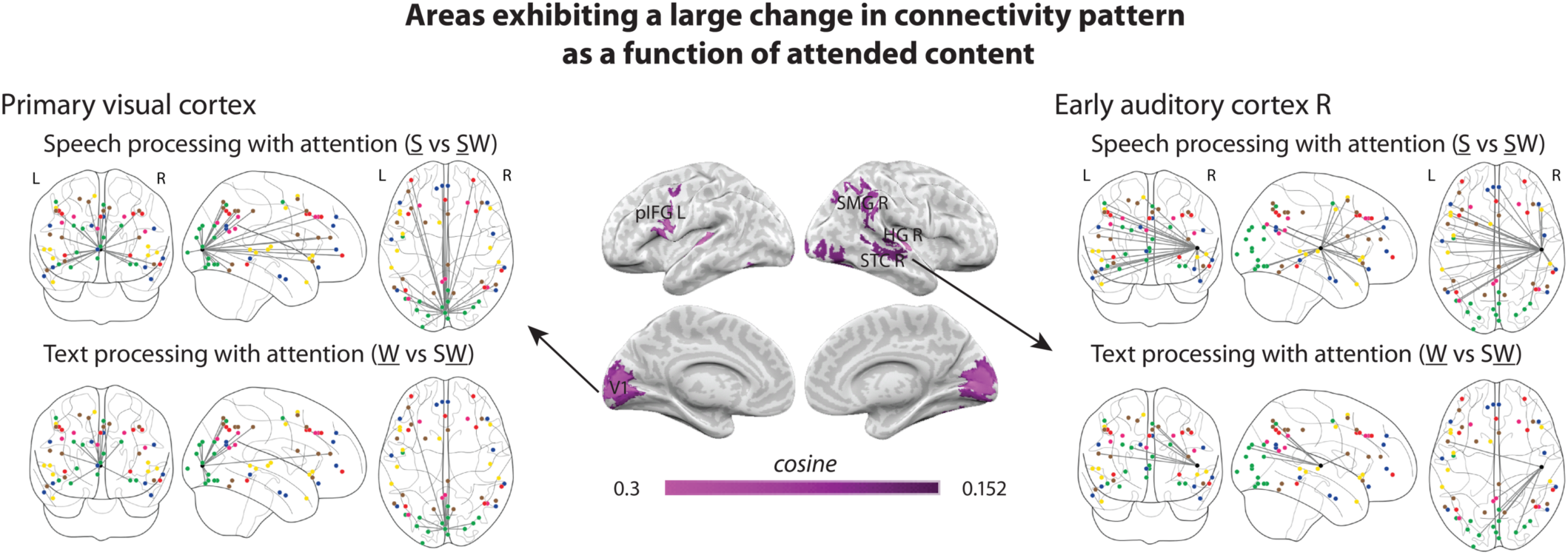
Regions that showed the largest change (top 20%) in their ISFC pattern as a function of whether the spoken or the written story was attended to. The change in ISFC pattern was assessed by calculating a cosine distance between a vector depicting coupling of goal-related written content (<u>W</u> subjects to S<u>W</u> subjects) and a vector depicting coupling of goal-related spoken content (<u>S</u> subjects to <u>S</u>W subjects). The strongest changes in shared spoken versus written content were observed in the primary visual and the right early auditory cortices.

**Table 2.** Hyper-induced topic search (HITS) scores and Cosine Distance scores of the 61 ROIs. The cell color of the score marks its strength relative to the group of regions (black: 1-2 SD, dark grey: 0-1 SD, light grey: 1-(−1) SD, white: −1-(−2) SD).

| Network | Area | HITS speech | HITS text | Cosine |
| --- | --- | --- | --- | --- |
| Visual | V1 | 0.011 | 0.0135 | 0.258 |
|  | mldV2 R | 0.0108 | 0.0139 | 0.193 |
|  | ldV2 L | 0.0105 | 0.0074 | 0.102 |
|  | vV2 | 0.0107 | 0.0143 | 0.203 |
|  | V3 | 0.0096 | 0.0124 | 0.104 |
|  | V3a R | 0.0109 | 0.0138 | 0.123 |
|  | V3a L | 0.0137 | 0.0153 | 0.143 |
|  | hV4 R | 0.0128 | 0.0143 | 0.023 |
|  | hV4 L | 0.0126 | 0.0108 | 0.112 |
|  | aCUN | 0.0118 | 0.0148 | 0.146 |
|  | dPCUN | 0.0136 | 0.0203 | 0.053 |
|  | LOC L | 0.014 | 0.0129 | 0.102 |
|  | LOC R | 0.0128 | 0.0139 | 0.169 |
|  | SOG L | 0.0087 | 0.013 | 0.139 |
|  | dPreCG L | 0.014 | 0.0079 | 0.195 |
| Auditory-Language | HG R | 0.0163 | 0.0068 | 0.908 |
|  | HG L | 0.0154 | 0.0069 | 0.877 |
|  | STC L | 0.0288 | 0.017 | 0.106 |
|  | STC R | 0.0263 | 0.0164 | 0.152 |
|  | SMA L | 0.0179 | 0.0126 | 0.149 |
|  | PMC L | 0.0199 | 0.0123 | 0.167 |
|  | aIFG L | 0.0225 | 0.018 | 0.11 |
|  | aIFG R | 0.0224 | 0.0171 | 0.075 |
|  | aSTG L | 0.0259 | 0.0187 | 0.066 |
| DMN-II | caMTG L | 0.0306 | 0.0245 | 0.053 |
|  | aMTG R | 0.0279 | 0.0208 | 0.063 |
|  | IOFG L | 0.0242 | 0.0186 | 0.102 |
|  | IOFG R | 0.0207 | 0.0167 | 0.073 |
|  | aANG R | 0.0211 | 0.0191 | 0.033 |
|  | aANG L | 0.0281 | 0.0245 | 0.048 |
|  | cMTG R | 0.0218 | 0.0187 | 0.147 |
|  | vmPFC | 0.0188 | 0.0147 | 0.091 |
|  | aPFC | 0.0246 | 0.0187 | 0.055 |
|  | SFG | 0.025 | 0.0192 | 0.065 |
| DMN-I | pANG R | 0.0203 | 0.0171 | 0.042 |
|  | pANG L | 0.0218 | 0.0166 | 0.097 |
|  | vPCUN | 0.0185 | 0.0181 | 0.095 |
|  | dPCC | 0.0153 | 0.0199 | 0.1 |
|  | dMFG R | 0.0109 | 0.0136 | 0.037 |
|  | dMFG L | 0.0137 | 0.0138 | 0.063 |
| Attention | pMTG R | 0.0137 | **0.0205** | 0.066 |
|  | IPL R | 0.0125 | **0.0212** | 0.106 |
|  | IPL L | 0.0149 | **0.0207** | 0.04 |
|  | dmPFG | 0.015 | 0.0185 | 0.051 |
|  | DLPFC L | 0.0194 | 0.019 | 0.096 |
|  | DLPFC R | 0.0136 | **0.0206** | 0.046 |
|  | aOFC L | 0.0138 | 0.0186 | 0.053 |
|  | aOFC R | 0.0128 | 0.0172 | 0.123 |
| Executive | pIFG L | 0.0174 | 0.0171 | 0.25 |
|  | vPFC R | 0.0139 | 0.0186 | 0.065 |
|  | smPFC | 0.013 | 0.0173 | 0.058 |
|  | dPoCS L | 0.01 | 0.019 | 0.068 |
|  | SFS L | 0.0089 | 0.0144 | 0.102 |
|  | dPoCS R | 0.0165 | **0.0203** | 0.125 |
|  | MCC | 0.0182 | 0.0168 | 0.139 |
|  | vMFG L | 0.0156 | 0.0197 | 0.044 |
|  | aINS L | 0.0117 | 0.0138 | 0.063 |
|  | SMG R | 0.015 | **0.0203** | 0.201 |
|  | SMG L | 0.011 | 0.0187 | 0.077 |
|  | pMTS R | 0.0095 | 0.0167 | 0.161 |
|  | pMTS L | 0.0106 | 0.0152 | 0.203 |

Second, we searched for central hubs that are connected to many other regions in each information matrix, and therefore are critical in each of the processing pathways. The degree of centrality of each area was assessed by considering the regions that are significantly correlated with it, using a hyperlink-induced topic search (HITS) algorithm. In undirected graphs like the thresholded ISFC matrix, this link analysis algorithm identifies a hub as a node linked to many other central nodes. Therefore, a region with high HITS score is not only highly connected, but also connected to other highly connected regions (see Materials and Methods). Each region’s degree of centrality was assessed separately for the attended spoken and attended written information graphs.

Most regions with a high degree of centrality were found in high-order multimodal areas of the brain, such as in the medial temporal, anterior frontal, and inferior parietal cortices (Fig. 10). The most highly connected hubs in both the spoken and written networks include caMTG L, the aMTG R, the aANG L, and the SFG. Furthermore, the degree of regions’ centrality was found to be quite stable across the two graphs. Areas with high (small) centrality while subjects attended spoken information also had high (small) centrality while subjects attended written information (*r* = 0.43; Fig. 10), thus demonstrating a certain degree of invariance to the identity of the story being processed.

**Figure 10.**
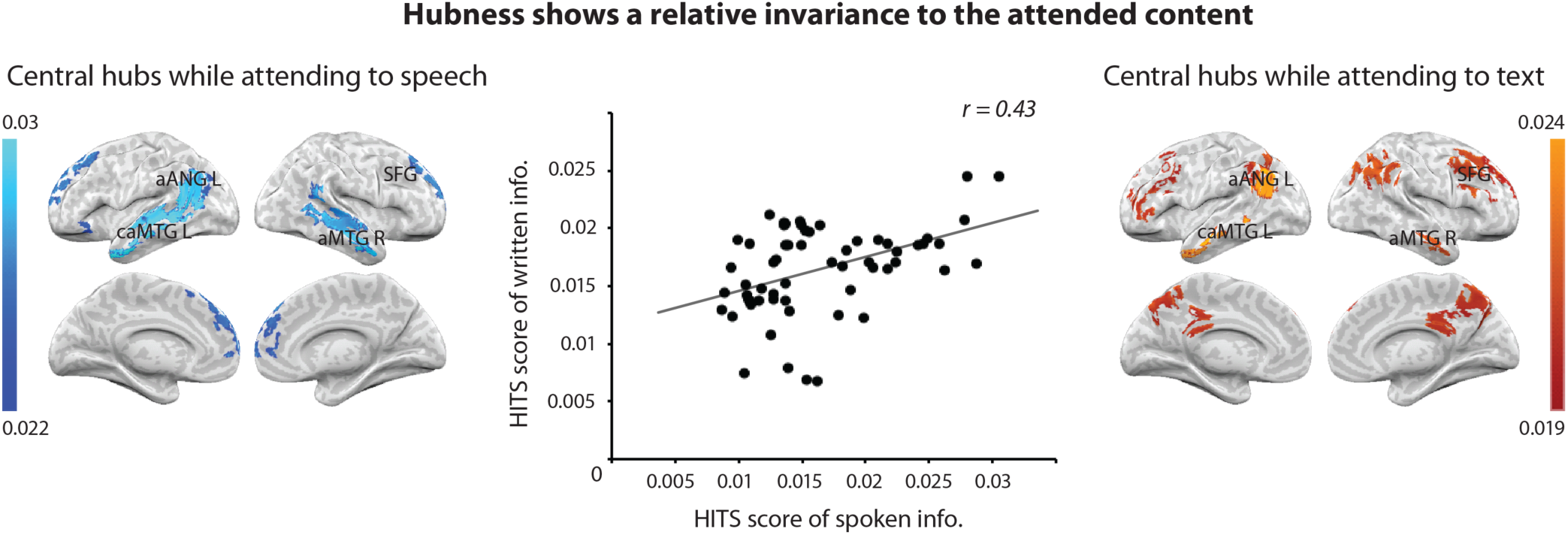
The degree of regions’ centrality (HITS) in the attended spoken information graph (<u>S</u> subjects to <u>S</u>W subjects) was correlated (*r* = 0.43) with their degree of centrality in the attended written information graph (<u>W</u> subjects to S<u>W</u> subjects). Regions that showed the highest (top 20%) degree of centrality in both attended spoken and written graphs included the caMTG L, the aMTG R, the aANG L, and the SFG.

Taken together, these results imply that by and large, early sensory regions alter their pattern of information routing as a function of the attended content, but they are less central in the processing networks of spoken or written information. At the same time, high-order brain regions tend to be relatively central in the processing pathways of both spoken and written stories, while the set of regions they are coupled with is relatively stable.

## Discussion

In this study, we tracked changes in the information shared across brain regions as a function of internal, attentional goals, while subjects were exposed simultaneously to two unrelated stories, one written and one spoken. A new approach for inter-subject functional correlation (ISFC), which compares the timecourses across cortical regions and conditions, enabled detection of changes in the way information from spoken and written stories was shared across the brain as a function of task. Specifically, written and spoken information was processed in sensory visual and auditory regions when it was attended and, to a lesser degree, when outside of the focus of attention (Fig. 5 and 7E). However, attention was required for the story-specific information to reach higher-order linguistic and extra-linguistic regions (Fig. 6). Furthermore, responses to the attended input stream not only propagate from sensory to higher-order brain regions along the processing hierarchy, but also reached intermediate sensory regions that processed the opposite unattended sensory information, perhaps reflecting a top-down influence or feedback from the higher-order areas (Fig. 8).

Our results show widespread attentional modulation along the processing hierarchy, with increasing selectivity for attended stories from early auditory and visual processing to high order linguistic and extra-linguistic areas. The processing of unattended information was weakly attenuated in sensory regions (Fig. 7), as was observed in previous neuroimaging studies (Watanabe et al., 1998; Gandhi et al., 1999; Petkov et al., 2004; Poghosyan and Ioannides, 2008). Such attenuation might reflect a diminished processing of the input in its earliest stages of perceptual processing, fitting the idea of early selection by attention (Hillyard et al., 1973; Posner and Driver, 1992; Woldorff et al., 1993). Alternatively, the modulation of response in sensory regions might result from selectivity in later stages of processing (e.g. semantic), which influence sensory regions through top-down feedback (Treisman, 1986; Wood and Cowan, 1995).

Regardless of the source of the attenuation in sensory regions, information from the irrelevant stimulus did not reach the majority of high-order cognitive regions that are typically involved in the processing of the story content when fully attended (Fig. 5). Indeed, we observed the greatest changes in the connectivity patterns as a function of attention in early sensory regions (Fig. 9), which supports the notion that attention operates at early stages of the processing hierarchy to select information to be further processed by higher-order brain regions. A similar hierarchical effect of attentional modulation in early sensory regions followed by a late strong selectivity to the attended speech in high-order regions was identified in a cocktail-party design using direct electrocorticography recordings (Golumbic et al., 2013). Here, we were able to demonstrate the effect in both the auditory and the visual systems during cross-modal competition, and expand the findings to inter-regional correlation patterns.

The limited spread of unattended information into high-order brain regions could be attributed to the diminished processing in sensory regions. Thus, the observed decline in response reliability in early sensory regions might reflect a filtering mechanism that prevents more complex processing in higher-order regions. For example, reduced signal reliability might reflect a decrease in the population’s SNR of the perceptual representation (Serences et al., 2009), or a lack of selective synchronization of neural oscillations that is crucial for further downstream processing (Stein and Sarnthein, 2000; Schroeder and Lakatos, 2009; Akam and Kullmann, 2010; Bosman et al., 2012).

In this study, we observed that signals related to the attended story also appeared into the secondary sensory regions of the competing *unattended* sensory modality (Fig. 8). Story-specific information from the attended *spoken* story was found in secondary *visual* regions, and story-specific information from the attended *written* story was found in secondary *auditory* regions. These intermediate sensory regions thus showed substantially mixed responses to both the attended and unattended stories (that were presented simultaneously). The presence of information from the attended story in the competing sensory pathway might indicate an attention based mechanism that prevents the spread of unattended information from sensory to higher-order brain regions (Baier et al., 2006; Mozolic et al., 2008). Alternatively, it could also reflect an excitatory signal that supports imagery evoked by the content of the attended story (Vetter et al., 2014). Further studies will be needed to determine the exact mechanism underlying the expanded neural response to attended content, and its influence on unattended sensory modalities.

We have demonstrated here for the first time that ISFC is a robust tool in tracking the changes in the information shared across brain regions as a function of top-down attention. By removing spontaneous neural responses, which contribute strongly to functional connectivity (FC), the sensitivity to stimulus-locked processes was improved (Simony et al., 2016), enabling the changes in propagation of information along the cortical hierarchy to be uncovered. ISFC could complement other methodological approaches to study the effects of top-down attention on inter-regional correlation. For example, several studies have regressed out stimulus-evoked responses and examined “background connectivity” in the residuals, which improved the sensitivity to intrinsic interactions between visual cortical regions of interest (Al-Aidroos et al., 2012; Norman-Haignere et al., 2012; Griffis et al., 2015; Cordova et al., 2016). ISFC examines the complementary side, capturing the stimulus-evoked interactions between regions while minimizing the intrinsic ones.

In conclusion, by following neural responses characteristic to each story, this study was able to show how intrinsic attention-based tasks can change the information shared along the processing hierarchy, from sensory regions to linguistic and extra-linguistic areas. We observed that attention modulated the processing of complex real-life narratives along both the visual and auditory pathways, with modulation increasing at successively higher levels of the cortical hierarchy, while unattended content was limited mainly to early sensory cortices. Furthermore, we detected the widespread reach of the attended content, which not only dominated most of the brain, but even appeared in secondary regions in the irrelevant sensory modality, perhaps reflecting top-down influence from higher-order brain regions. These findings improve our understanding of the complex influence of attentional control on the processing of spoken and written language in a multimodal environment, and open the door for future exploration of how information is dynamically routed between brain areas.

## Acknowledgments

This work was supported by the National Institute of Health (grant number DP1HD091948-01 to U.H. and M.R.) and the William Orr Dingwall Foundation (to M.R.). We thank Hanna Hillman for editing the manuscript and Chris Honey, Chris Baldassano, Nick Turk-Browne and the members of the Hasson lab for their helpful comments.

## Materials and Methods

### Subjects

Seventy-four subjects successfully participated in one of the two main experimental conditions (attention to text and attention to speech), or in one or two of the control conditions (unimodal text and speech). Eighteen subjects were discarded from the analysis: 6 subjects due to head motions >3mm, 5 for closing their eyes during the story, 1 due to corrupted anatomical signal, 1 due to anomalous anatomy, 1 due to difficulties in hearing the stimulus, 1 due to missing behavioral results, 1 due to previous familiarity with the story, and 2 due to failure in the memory tests (see behavioral assessment). Additional subjects were scanned until data from 18 subjects were collected for the attention to text (14 females; ages 19-32), attention to speech (14 females; ages 18-24), unimodal text (13 females; ages 18-29), and unimodal speech (13 females; ages 18-30) conditions. Sixteen of the subjects participated in both the unimodal text and speech conditions. Another 19 subjects were scanned in a rest condition (8 females, ages 18-31), and another 13 subjects were scanned while presented with a different audiovisual story (7 females, ages 19-26).

Procedures were approved by the Princeton University Committee on Activities Involving Human Subjects. All subjects were right-handed native English speakers, reported normal hearing and normal or corrected-to-normal vision, normal reading skills, and had not heard the two stories prior to the experiment. All subjects provided written informed consent.

### MRI acquisition

Subjects were scanned in a 3T full-body MRI scanner (Skyra, Siemens) with a 20-channel head coil. For functional scans, images were acquired using a T2*-weighted echo planer imaging (EPI) pulse sequence [repetition time (TR), 1500 ms; echo time (TE), 28 ms; flip angle, 64°], each volume comprising 27 slices of 4 mm thickness with 0 mm gap; slice acquisition order was interleaved. In-plane resolution was 3×3 mm^2^ [field of view (FOV), 192×192 mm^2^]. Anatomical images were acquired using a T1-weighted magnetization-prepared rapid-acquisition gradient echo (MPRAGE) pulse sequence (TR, 2300 ms; TE, 3.08 ms; flip angle 9°; 0.89 mm^3^ resolution; FOV, 256 mm^2^). To minimize head movement, subjects’ heads were stabilized with foam padding.

Subjects were provided with an MRI compatible in-ear mono earbuds (Sensimetrics model S14), which provided the same audio input to each ear. MRI-safe passive noise-canceling headphones were placed over the earbuds for noise removal and safety. The text was projected using an LCD projector onto a rear-projection screen located in the magnet bore, and was viewed with an angled mirror. Stimuli were presented and synchronized with MRI data acquisition onset using the Psychophysics toolbox (Brainard, 1997; Pelli, 1997) for MATLAB.

### Stimuli

The spoken language stimulus was a 15:03 min real-life story (“Slumlord” told by Jack Hitt, recorded live at “The Moth” storytelling event, New York City). The written language stimulus was a 15:03 min transcript of a different real-life story (“The Overview Effect” told by Richard Garriott), recorded at the same live storytelling performance. The content of the two stories was not related.

In the written stimulus, the words were individually presented in a white font in the center of a black screen in a rapid serial visual presentation. Out of the 2381 words of the story, most of the words (2093) were presented for the duration of 310 ms each. 189 of the resulting words were accompanied with punctuation and thus presented for 650 ms, and the remaining 99 words appeared at sentences’ ends and thus presented for 1200 ms. These varied durations aimed at providing an easy reading experience for subjects (Castelhano and Muter, 2001). The width of the words ranged from 4 to 160 pixels, the height ranged from 11 to 20 pixels.

In addition to the written and/or spoken stimulus, all conditions contained a red fixation point (radius of 5.6 pixels, 30% transparency) at the center of the screen, juxtaposed over the center of the words of the written stimulus, if presented. Neutral lead-in music was played for 12 s before the onset of the spoken stimulus, and graphical music symbols were shown for 12 s before the onset of the written stimulus. Responses to these initial 12 s were excluded from all analyses.

The spoken and written stories were combined to create a simultaneous auditory and visual presentation in the two main experimental conditions (Fig. 1). In the two control conditions, either the spoken or written stories were presented on their own, in a unimodal fashion

### Experimental design

Participants were instructed to carefully attend and remember the details of the presented story and were informed they would receive a monetary bonus based on their performance in the subsequent memory test. In the two main conditions where the two stories were presented simultaneously, participants were instructed to ignore the distracting story: The attention to text group (S<u>W</u>) attended the written stimulus while ignoring the spoken one, while the attention to speech group (<u>S</u>W) did the opposite (Fig. 1). In the two additional control groups, participants were exposed to one of the stories alone: The unimodal text group (<u>W</u>) read the written story, and the unimodal speech group listened to the spoken story (<u>S</u>).

All participants who were to be exposed to written content in the experiment got to practice reading a sample of unrelated text presented in RSVP before the beginning of the trial. The volume of the auditory stimuli was adjusted individually for each subject to a comfortable and clear level. When the spoken stimulus was attended in the attention to speech and unimodal speech groups, subjects were asked to fix their gaze toward the fixation point for the entire time. In the cases where subjects participated in both of the control conditions (89% of the subjects), the order of the two conditions was randomized.

### Data analysis

#### Preprocessing

fMRI data was preprocessed in FSL (http://fsl.fmrib.ox.ac.uk/fsl), including slice time correction, motion correction, linear detrending, high-pass filtering (140 S cutoff), and coregistration and affine transformation of the functional volumes to a template brain (MNI). Functional images were resampled to 3 mm isotropic voxels for all analyses. All calculations were performed in volume space. Projections onto a cortical surface for visualization were performed, as a final step, with NeuroElf (http://neuroelf.net).

#### Inter-subject Functional Correlation (ISFC) analysis

We calculated the ISFC matrix between all ROIs across brains (1) of subjects within the same condition (Simony et al., 2016) and (2) of subjects from two different conditions (Fig. 3). The neural signals *X*_*i*_ measured from subject *i*, *i* = 1, …, *k* are in the form of a *p*×*n* matrix that contains signals from *p* neural sources over *n* time points. All timecourses were z-scored within subjects to zero mean and unit variance. Thus, the subject-based ISFC was calculated by the Pearson correlation between single subject and the average of all other subjects as:

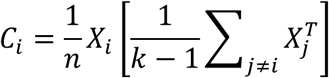

Hence, the *p*×*p* group-based ISFC matrix was given by:

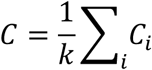

Subject-based ISFC between two conditions was calculated by the Pearson correlation between single subject from one group and the average of all subjects from the other condition as:

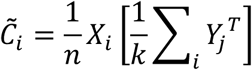

where the neural signals *Y*_*i*_ measured from subject *i*, *i* = 1, …, *k*. The cross-group ISFC matrix was given by:

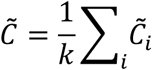

The final ISFC matrix is given by (*C* + *C*^*T*^)/2 within a group. This symmetry was imposed because the correlation between two brain regions was considered to be undirectional, as in FC. Similarly, the final ISFC matrix between two groups is given 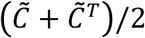, and then averaged with the final ISFC matrix calculated between individuals from the second group and averaged signal from the first group.

The ISFC was performed across the following conditions: attention to speech condition vs. unimodal speech control condition, attention to text condition vs. unimodal text control condition, attention to text condition vs. unimodal speech control condition, attention to speech condition vs. unimodal text control condition, and unimodal text control condition vs. unimodal speech control condition. ISFC was also performed within each of the four groups (see Supplementary Fig. 1A and C for groups <u>S</u> and <u>W</u>).

#### Inter-subject Correlation analysis

ISC maps were produced across conditions (e.g., attention to speech group vs. unimodal speech control group). The ISC maps provide a measure of the similarity of brain responses between two different conditions by quantifying the correlation of the timecourse of BOLD activity between each subject from one group and the averaged activity of all subjects in the other group (Hasson et al., 2004; Honey et al., 2012).

For each voxel, ISC between two conditions is calculated as an average correlation:

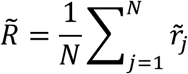

where the individual 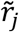 are the Pearson correlations of that voxel’s BOLD timecourses of the *j*’th individual from the first group and the average of that voxel’s BOLD timecourse of all individuals in the other group (Fig. 3).

The ISC was performed across the following conditions: attention to speech group vs. unimodal speech group; attention to text group vs. unimodal text group; attention to text group vs. unimodal speech group; attention to speech group vs. unimodal text group; unimodal text group vs. unimodal speech group.

ISC within a condition is calculated as an average correlation:

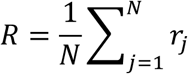

where the individual *r*_*j*_ are the correlations between the BOLD timecourse in one individual and the average of the BOLD timecourses in the remaining individuals. This within condition ISC analysis was performed within the group presented with a coordinated audiovisual story (see ROIs analysis).

In a standard GLM analysis, experimenters usually assume a prototypical response profile for each specific stimulus. The ISC analysis method differs from conventional fMRI data analysis methods in that it circumvents the need to specify a model for the neuronal processes for any given condition. Instead, when performed across conditions, the ISC method uses the averaged brain response to the content in one condition within a given brain area as a model to predict brain responses to different content presented in the other condition. When performed within a condition, the ISC method uses the subject’s brain responses as a model to predict brain responses to the same content.

#### ISFC and ISC bootstrapping and phase-randomization

Because of the presence of long-range temporal autocorrelation in the BOLD signal (Zarahn et al., 1997), the statistical likelihood of each observed correlation was assessed using a bootstrapping procedure based on phase randomization. The null hypothesis was that the BOLD signal in each area in each individual was independent of the BOLD signal values in the corresponding area in any other individual at any point in time (i.e., that there was no ISFC or ISC between any pair of subjects).

For all conditions, a phase randomization of each voxel timecourse was performed by applying a fast Fourier transform to the signal, randomizing the phase of each Fourier component, and inverting the Fourier transformation. This procedure scrambles the phase of the BOLD timecourse but leaves its power spectrum intact. For each randomly phase-scrambled surrogate dataset, we computed the ISC or ISFC (R) for all areas in the exact same manner as the empirical cross-group correlation maps described above. I.e. for ISC, by calculating the Pearson correlation between that voxel’s BOLD timecourse in one individual from one group and the average of that voxel’s BOLD timecourses of all individuals from the other group. For ISFC, by calculating the Pearson correlation between a region’s BOLD timecourse in one individual and the average of another region’s BOLD timecourse of all individuals from the other group. The resulting correlation values were averaged within each voxel (for ISC) or each pair of regions (for ISFC) across all subjects, creating a null distribution of average correlation values for all voxels or all pair of regions.

To correct for multiple-comparisons, we selected the highest ISC (or ISFC) value from the null distribution of all voxels (or pair of regions) in a given iteration. We repeated this bootstrap procedure 10,000 times to obtain a null distribution of the maximum noise correlation values (i.e., the chance level of receiving high correlation values across all voxels in each iteration).

Because the participants in the two unimodal groups (<u>S</u> and <u>W</u>) were exposed to two different external stimuli with no common features (i.e., distinct modality and content), we assumed that the inter-subject correlation between these groups could be regarded as noise. Thus, the probability of the highest and lowest ISC (or ISFC) correlation value measured between these two groups (highest: 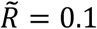, 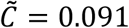; lowest: 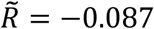, 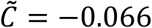) was assed based on the null distribution of the maximal noise correlation values (highest: *p* = 1.56×10^−44^, *p* = 3.8×10^−28^; lowest: *p* = 3.8×10^−28^, *p* = 3.8×10^−28^). Next, inter- or intra-regional correlations (ISFC or ISC) were retained if their probability exceeded the probability of the maximum correlation value (or fell below the probability of the minimum correlation value) measured between the two unimodal groups. Specifically, familywise error rate (FWER) was defined for each of the null distributions of the different cross-group comparisons (<u>S</u> vs. <u>S</u>W, <u>S</u> vs. S<u>W</u>, <u>W</u> vs. S<u>W</u>, <u>W</u> vs. <u>S</u>W) according to the probability of the maximal/minimal correlation value measured between the unimodal groups. The corresponding correlation values in each cross-group null distributions (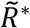 or 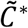) were used to threshold the veridical cross-group correlation data (Nichols and Holmes, 2001). In other words, in the ISC map, only voxels with mean correlation value 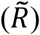 above the threshold derived from the bootstrapping procedure 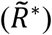 were considered significant after correction for multiple-comparisons and were presented on the final map. In the ISFC graph, only pairs of regions with mean correlation value 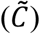 above the threshold derived from the bootstrapping procedure 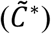 were considered significant after correction for multiple-comparisons and were presented on the final map.

Using this method, the thresholds for each cross-group comparison were as follows: Attention to speech group vs. unimodal speech group 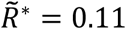 and 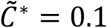; attention to text group vs. unimodal text group 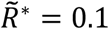 and 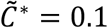; attention to text group vs. unimodal speech group 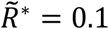 and 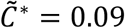; attention to speech group vs. unimodal text group 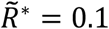 and 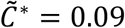.

The same procedure was performed on the within-group correlation maps (ISC), in which correlations were computed between each individual and the average of the remaining participants in the group, as described above. In these cases, the FWER was defined as the top 0.1% of the null distributions (instead of based on the comparison between the unimodal groups). Using this method, the thresholds for within-group comparisons were as follows: Unimodal speech group (<u>S</u>) *R*^∗^ = 0.097 (see Supplementary Fig. 1A and B); unimodal text group (<u>W</u>) *R*^∗^ = 0.098 (see Supplementary Fig. 1C and D); The group of participants presented with the coordinate audiovisual story *R*^∗^ = 0.15 (see ROI analysis).

#### Attention-Index (AI)

To measure effects of attention on processing speech, a modulation index was calculated for both ISC and ISFC within each brain area as (|*corr*(*<u>S</u>W*,*<u>S</u>*)| − |*corr*(*S<u>W</u>*,*<u>S</u>*)|)/(|*corr*(*<u>S</u>W*,*<u>S</u>*)| + |*corr*(*S<u>W</u>*,*<u>S</u>*)|); to measure effects on processing text the AI was calculated as (|*corr*(*S<u>W</u>*,*<u>W</u>*) − |*corr*(*<u>S</u>W*,*<u>W</u>*)|)/(|*corr*(*<u>S</u>W*,*<u>W</u>*)| + |*corr*(*<u>S</u>W*,*<u>W</u>*)|). This analysis was applied only to regions that show reliable responses to the story in either its attended or unattended state (i.e., noisy regions that did not respond to the stories were removed from the analysis; see Materials and Methods, under *ISFC and ISC bootstrapping and phase randomization*).

#### Segregation-Index (SI)

To measure how segregate are the responses to the attended speech and unattended text presented simultaneously in condition <u>S</u>W, a segregation index was calculated for both ISC and ISFC within each brain area as: (|*corr*(*<u>S</u>W*, *<u>S</u>*)| − |*corr*(*<u>S</u>W*,*<u>W</u>*)|)/(|*corr*(*<u>S</u>W*, *<u>S</u>*)| + |*corr*(*<u>S</u>W*,*<u>W</u>*)|); To measure how segregated are the responses to the attended text and unattended speech presented simultaneously in condition S<u>W</u>, the SI was calculated as (|*corr*(*S<u>W</u>*, *<u>W</u>*)| − |*corr*(*S<u>W</u>*,*<u>S</u>*)|)/(|*corr*(*S<u>W</u>*, *<u>W</u>*)| + |*corr*(*S<u>W</u>*,*<u>S</u>*)|),. This analysis was applied only to regions that show reliable responses to the story in either its attended or unattended state (i.e., noisy regions that did not respond to the stories were removed from the analysis; see Materials and Methods, under *ISFC and ISC bootstrapping and phase randomization*).

#### Partial correlation

Under the Gaussian assumption, it is known that the partial correlation (conditional dependency) is characterized by the inverse covariance matrix (Θ). Thus, an undirected graph encoding conditional dependencies among the ROIs can be constructed based on the inverse covariance matrix. In particular, we place an edge between ROIs *i* and *j* if and only if Θ_*i*,*j*_ ≠ 0. To estimate sparse graphs, we used a constrained l1-minimization approach for estimating the inverse covariance matrix (Cai et al 2011, Supplementary Fig. 4). By adding an l1 penalty in the minimization problem, it encourages the elements in the estimated inverse covariance matrix to contain many zero elements. This algorithm has the advantage over others by not requiring the covariance matrix to be symmetric positive semidefinite.

#### Degree of Centrality analysis

Each region’s degree of centrality was assessed separately for the attended spoken and attended written information graphs. We performed a hyper-induced topic search (HITS) algorithm on the two ISFC graphs of <u>S</u> vs. <u>S</u>W and <u>W</u> vs. S<u>W</u> (Table 2). An ROI with a high HITS score is highly connected to other regions in the graph, that are themselves highly connected. We identified the most highly connected regions within each of the networks as those with the top 20% HITS scores (Fig. 10). To assess the similarity in regions’ relative centrality in each functional network, Pearson correlation was performed over all HITS scores across the two graphs.

#### Vector Similarity analysis

We examined how pattern of correlations changed as a function of the input attended for each ROI. The pattern of ISFC in each of the 61 ROIs could be considered as a point in a 60-dimensional space, or a vector whose angle and length can be defined relative to the origin (Wang et al., 2015). We defined these correlation vectors for <u>S</u> vs. <u>S</u>W and <u>W</u> vs. S<u>W</u>, which resulted in two correlation vectors for each ROI. We extracted the angle change (cosine distance) between the two correlation vectors, with large distance between the vectors capturing a large change in goal-related inter-regional coupling patterns (Table 2). We identified the regions that showed the largest change in their ISFC pattern as those with the top 20% cosine distance (Fig. 9).

#### Functional Connectivity (FC) analysis

The FC correlation matrix was calculated between all voxels within the brains of subjects from the resting group. The neural signals *X*_*i*_ measured from subject *i*, *i* = 1, …, *k* are in the form of a *p*×*n* matrix that contains signals from *p* neural sources over *n* time points. All timecourses were z-scored within subjects to zero mean and unit variance. Thus, the subject-based FC was calculated by the Pearson correlation between different voxels within a single subject and then averaged across all group members.

### Intrinsic connectivity networks (ICNs) and Regions of Interest (ROI) analysis

First, we excluded voxels that were not demonstrated before as showing a reliable response to written and spoken content. For this purpose, voxel-based ISC analysis was performed over a separate group of subjects exposed to a coordinated audiovisual presentation of the same story (“Pie-man”, see Regev et al. 2013 for more stimulus related details). In the process of defining ROIs and intrinsic connectivity networks (ICNs), only voxels that demonstrated a significantly reliable response to the coordinated audiovisual linguistic stimulus were included.

Next, we mapped ICNs by clustering patterns of functional connectivity that were calculated in a separate group of resting participants. FC analysis was performed between all filtered voxels within each of the 19 subjects and then averaged across all group members. K-mean clustering was performed over the FC correlations using kmeans function in MATLAB to extract six ICN. This procedure partitioned the voxel-based FC correlation matrix into k mutually exclusive clusters. Each cluster was defined by a set of N member voxels (each with an associated correlation vector) and by the centroid of the correlation vectors in the cluster. The iterative algorithm minimizes the sum of distances from each voxel (vector) to its cluster centroid, over all clusters. Each cluster contained functionally connected voxels that were grouped as a network. The extracted networks were labeled based on their anatomical identification as: Auditory, Visual, DMN-I, DMN-II, Attention, and Executive (Supplementary Fig. 2).

The six ICNs were then split into regions of interest (ROIs). We started by identifying the early auditory and sensory cortices within the auditory and visual networks. Heschl’s gyri were defined within the auditory network based on the probabilistic Harvard-Oxford Cortical Structural Atlas (Desikan et al., 2006). V1, V2, V3, V3a and hV4 were identified within the visual network using a published retinotopic probabilistic atlas (Wang and He, 2014). In cases where a voxel was included in several regions simultaneously, it was assigned to the region with the highest probability to avoid overlapping ROIs. Four voxels from V1 that did not overlap with the visual network were discarded.

In further defining ROIs in the rest of the networks, voxels were clustered using a parcellation approach based on local connectivity patterns (Baldassano et al., 2015). This clustering method was performed within each network on the FC voxelwise correlations of the resting group, excluding the early sensory regions defined earlier. Each network was clustered into the minimal number of exclusive regions that would not extend 900 voxels each.

Finally, regions that were not spatially continuous (e.g., across hemispheres) were split. Along the process of defining ROIs, any region that included less than 10 voxels was dropped from further analysis, adding up to 271 dropped voxels in total.

Overall, this procedure yielded 61 ROIs: 15 in the visual network, 9 in the auditory network, 10 in the DMN-II, 6 in the DMN-I, 8 in the attention network, and 13 in the Executive network (Table 1; Fig. 4).

### Network visualization

The pattern of ISFC for two selected ROIs with the rest of the brain (Fig. 9) was represented on a schematic brain with 61 nodes representing the ROIs. Each node was located at the centroid of the ROI, and each significant correlation was described as a line between the nodes.

### Behavioral assessment

Immediately following the scan, each subject’s comprehension, memory and engagement for the two stories was assessed using computerized questionnaires. Subjects in the simultaneous conditions were informed in advance about a memory test only for the story they were asked to attend, but were surprised after the scan with an additional memory test for the unattended story. Before answering the tests, it was explained to them that correct responses would win them a monetary bonus, regardless of the story. This procedure aimed at encouraging participants to answer the memory tests to the best of their abilities, regardless of whether they were originally instructed to attend them or not.

The order of the two story questionnaires was counterbalanced between subjects within each of the conditions. Subjects in the control conditions were also informed in advance of the memory tests for the one or two unimodal stories they attended (no unattended content presented). For the subjects who participated in both control conditions and therefore attended (separately) both stories, the order of the two questionnaires was pseudo-randomized within each of the conditions. The order of the questions within each story questionnaire was not randomized.

#### Free recall

For the first memory test, subjects were asked to write down a summary of the narrative of the story, as detailed as possible. Three independent raters graded these written records (on a scale from 1 to 20), taking into account subjects’ general story comprehension as well as memory for small details. Raters’ grades were highly cohesive. The scores were z-scored within each rater, averaged across all three of them, and linearly transformed to positive values. One subject’s recall summary was not included in the analyses due to a technical problem in saving the data.

#### Multiple choice

In the second memory and comprehension test, subjects were asked to answer forced-choice questions with four potential choices about the content of the story (24 questions for the written story and 25 for the spoken story). The percentage of correct answers was calculated for each subject.

#### Fill-in-the-blank

For the last memory test, subjects were given the story’s transcript with missing words or phrases (84 in the written story and 77 in the spoken story) and were asked to fill in the missing text. Importantly, no information related to the missing words appeared in the previous multiple-choice questions. Three independent raters graded each of the fill-ins (on a scale from 0 to 4), taking into account the subjects’ general conceptual memory for the missing text as well as the accuracy of the wording. The percentage of the score received out of the total maximal score was calculated for each of the subjects and averaged across all three raters.

For each story questionnaire, independent sample t-tests were conducted to compare the effect of the task on (a) the quality of the rated free-recall test; (b) the success in the multiple-choice test; and (c) the success in fill-in-the-blank memory tests. The tests were calculated between the attention to text group (S<u>W</u>) and the attention to speech group (<u>S</u>W), between the attention to text group and the unimodal text group (<u>W</u>), and between the attention to speech group and the unimodal speech group (<u>S</u>). Effect sizes were assessed using Cohen’s *d*.

An exclusion criterion was applied based on participants’ multiple-choice memory and comprehension performances to screen for individuals who did not perform their tasks as expected. According to this criterion, any participants who deviated from the minimum requirement of 65% correct responses for the attended stimuli were excluded from both the behavioral and neural analysis. Two participants from the group attending the spoken story were marked as outliers based on their low performance in the spoken story multiple choice test.

### Eye tracking

Eye tracking was conducted using the iView X MRI-LR system (Sensomotoric Instruments [SMI]), sampling at 60Hz. The experimenter monitored participants’ alertness and general direction of gaze via the eye tracking camera. Any participant who appeared not to be looking toward the monitor or who closed their eyes was excluded from all analyses. In addition, some participants were not included in the eye tracking analysis due to calibration problems or incomplete eye tracking data (i.e., missing more than 60% of the data), including 3 subjects from the attention to speech condition, 2 subjects from the attention to text condition, and 2 subjects from the unimodal control conditions.

Eye movements were measured to examine deviations in gaze from the written stimulus. For this purpose, the proportion of the gaze data that remained within the horizontal borders of the longest written word of the story (160 pixels long) was calculated for each subject. All participants remained within these borders for 95% of the recorded data samples, except for one participant in the attention to text condition (92% of data within the borders) and one participant in the unimodal speech control (91% of the data within the borders).

## Supplementary Material

**Supplementary Figure 1.**
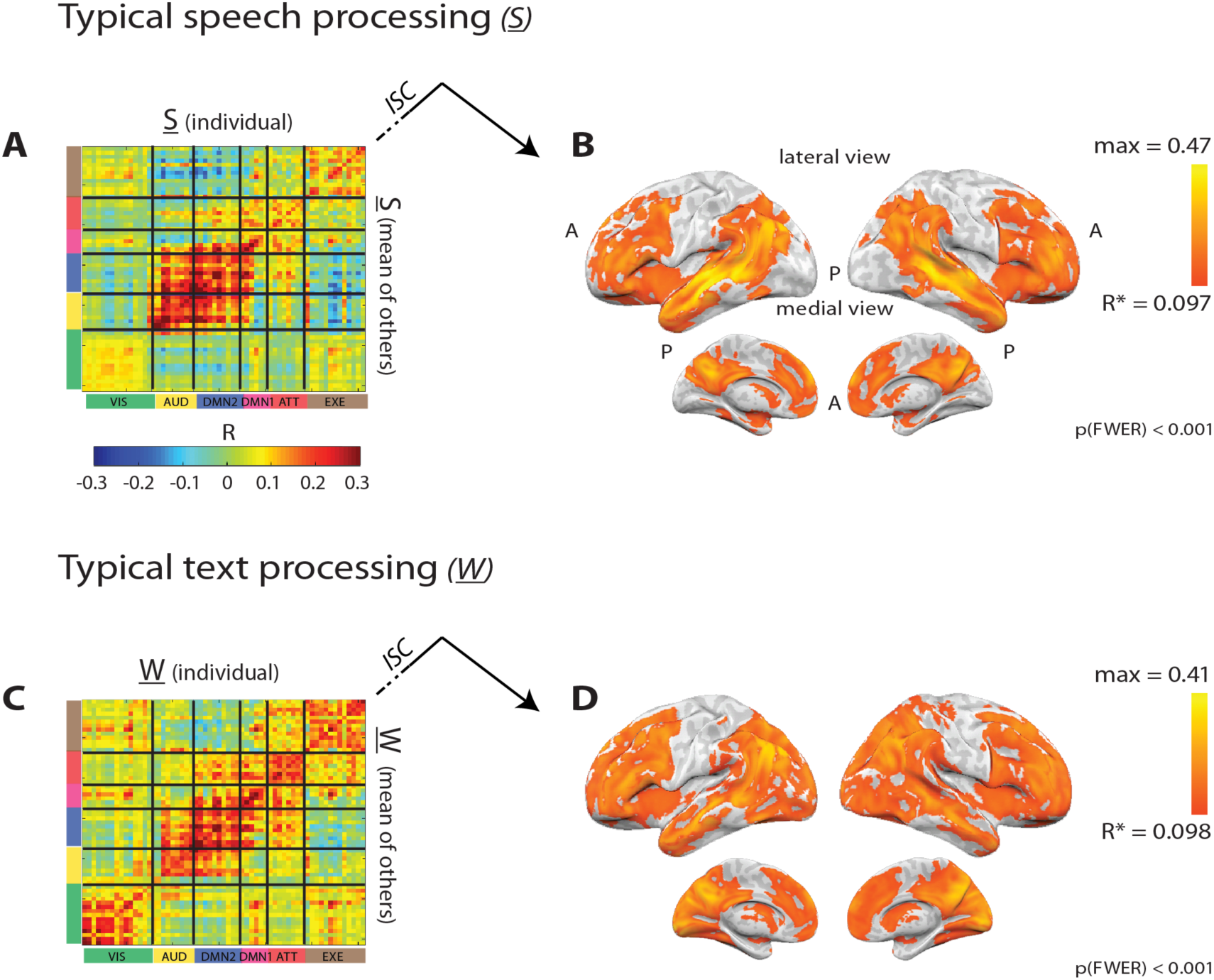
Reliable inter- and intra-regional responses to spoken (**A** and **B**) and written (**C** and **D**) stories. The BOLD timecourse was correlated within the same brain area (ISC) or between different brain areas (ISFC) across subjects within the same unimodal control conditions (<u>S</u> or <u>W</u>).

**Supplementary Figure 2.**
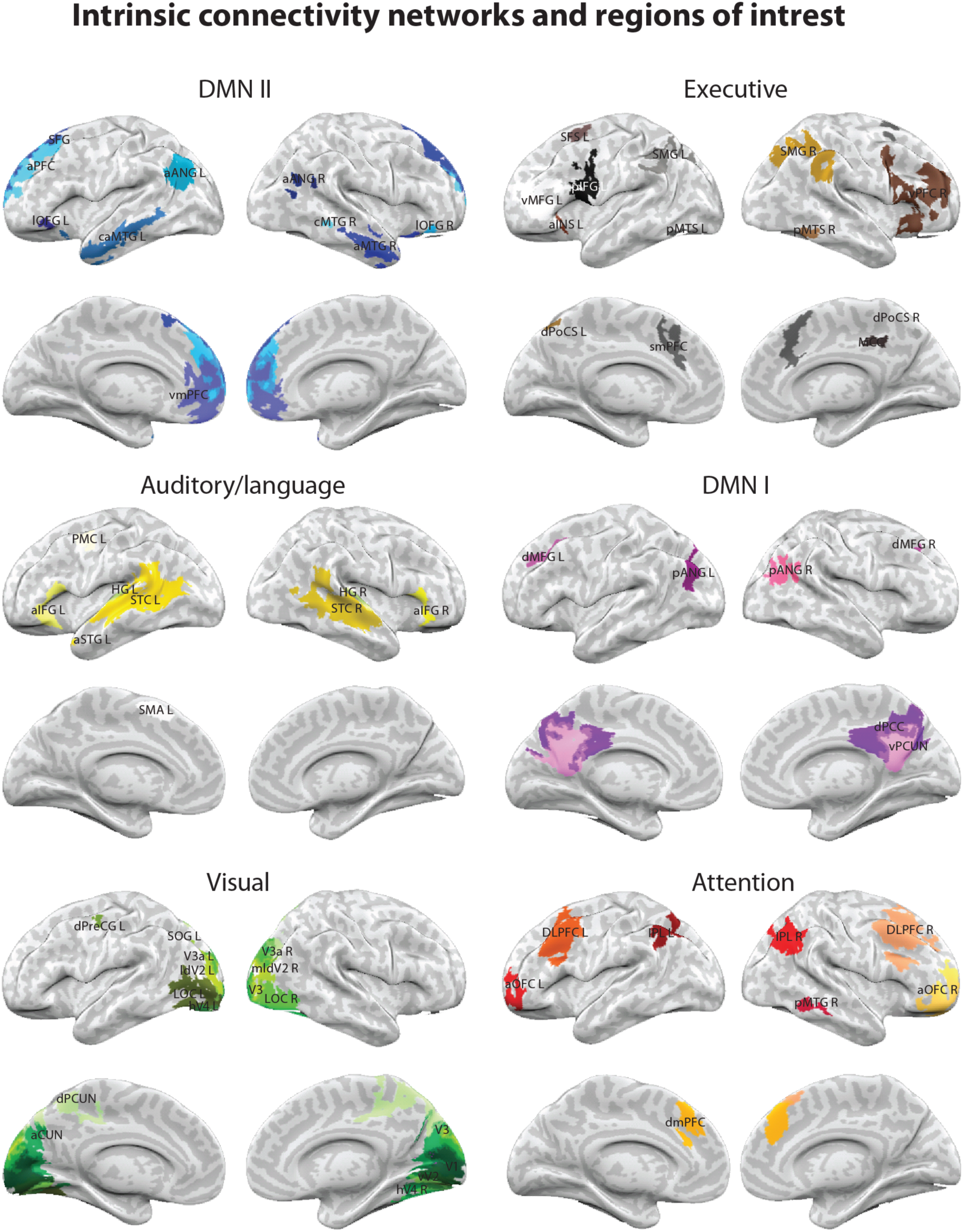
Six intrinsic connectivity networks (ICNs), split into 61 regions of interest (ROIs). For more information about the definition process of the different networks and regions see Materials and Methods.

**Supplementary Figure 3.**
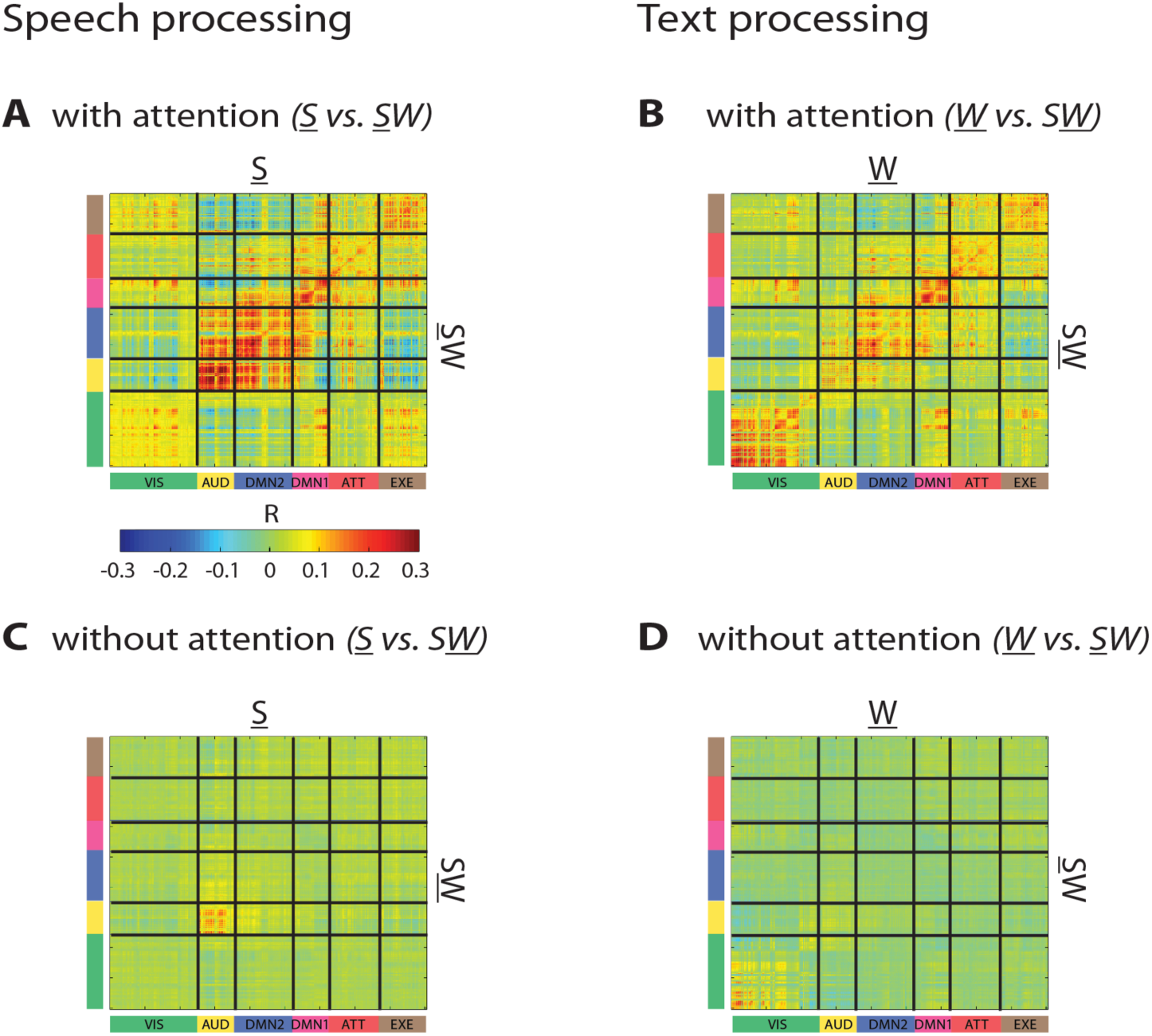
Voxel-based ISFC analysis across groups showed that attention to spoken (**A**) and written (**B**) stories evoked typical processing responses that spread from sensory areas into high-order areas. However, the typical response to the spoken (**C**) and written (**D**) stories were limited to the auditory and visual cortices respectively when ignored.

**Supplementary Figure 4.**
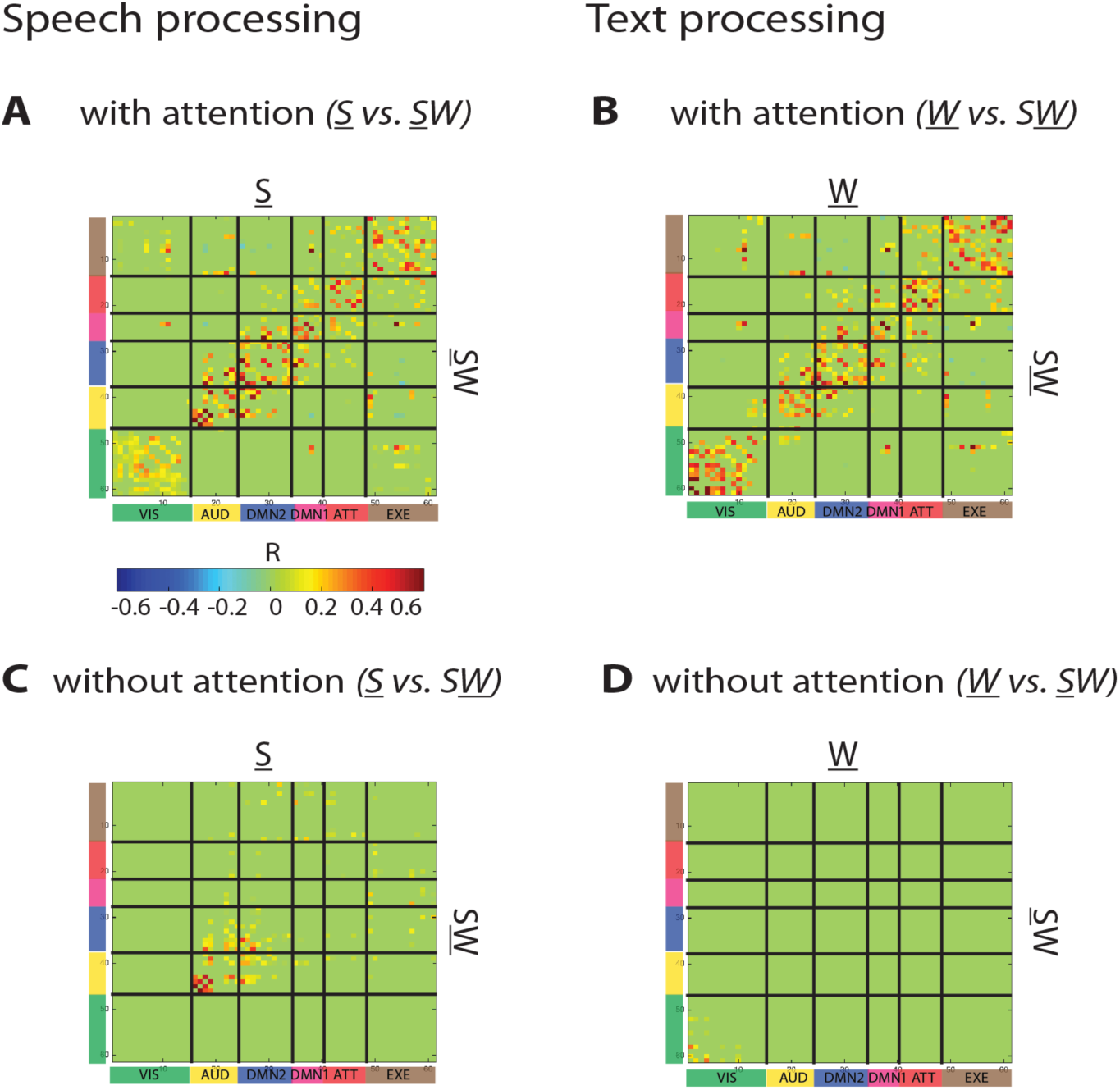
The mutual dependencies between ROIs were removed when ISFC was performed across groups, to reflect more direct inter-regional interaction (i.e. using a partial correlation analysis). Attention to spoken (**A**) and written (**B**) stories evoked typical processing responses that spread from sensory areas into high-order areas. However, the typical responses to the spoken (**C**) and written (**D**) stories were limited to the auditory and visual cortices correspondingly when ignored.

